# B cell-intrinsic IRF8 transcriptionally reprograms antigen presentation to sustain CD8⁺ T cell antitumor immunity

**DOI:** 10.64898/2026.06.28.735129

**Authors:** Zainab Tiamiyu, Dakota B. Poschel, Richa Rashmi, Sergei Bombin, Kendra Fick, Patrick Czabala, Dafeng Yang, Huidong Shi, Keita Saeki, Keiko Ozato, Kebin Liu

## Abstract

Interferon regulatory factor 8 (IRF8) is a master transcription factor of myeloid differentiation, but whether IRF8 intrinsically controls B cell function in tumors remains unknown. Using paired single-cell transcriptomic and chromatin accessibility profiling of tumors from wild-type and Irf8-deficient mice, we identify a B cell-intrinsic IRF8 axis regulating antigen presentation and sustaining anti-tumor CD8⁺ T cell immunity. IRF8 establishes conserved chromatin accessibility programs across myeloid cells and plasmablasts centered on antigen processing and MHC class I presentation, but engages distinct motifs by lineage: myeloid cells preferentially utilize ISRE and ETS-composite elements, whereas plasmablasts are selectively enriched for EICE elements, reflecting B lineage-specific IRF8-IRF4 cooperation. Loss of IRF8 disrupts these programs, skews B cells toward plasmablast differentiation and reduces antigen presentation machinery. B cell depletion accelerated tumor growth, while CD40 agonism activated B cells, expanded T cells, and enhanced anti-tumor immunity. B cell-specific IRF8 deletion alone accelerated tumor growth, establishing a cell-intrinsic requirement independent of myeloid IRF8 function. The IRF8-regulated B cell signature was enriched in PD-1 blockade cancer patient responders, and plasmablast abundance correlated with response in pembrolizumab-treated cancer patients. These findings establish IRF8 as a lineage-adapted regulator of antigen presentation and define the IRF8-B cell axis as a determinant of anti-tumor immunity.

**Highlights:** IRF8 establishes a conserved chromatin accessibility across tumor-infiltrating myeloid and B cells

Myeloid cells engage ISRE motifs, whereas plasmablasts rely on EICE motifs as IRF8 lineage-specific cis-regulation

IRF8 regulates an antigen presentation in B cells to sustain anti-tumor T cell immunity

B cell-intrinsic IRF8 transcription signature predicts patient response to PD-1 blockade immunotherapy

## INTRODUCTION

Interferon regulatory factor 8 (IRF8) is a member of the IRF transcription factor family that directs bipotential myeloid progenitors toward macrophage and dendritic cell (DC) fates while restraining granulocytic differentiation ^1–3^. In both mice and humans, IRF8 deficiency causes profound defects in conventional type 1 dendritic cell (cDC1) and plasmacytoid DC (pDC) development, underscoring its conserved role in antigen-presenting cell (APC) specification ^4,5^. IRF8 maintains cDC1 identity through preservation of lineage-specific chromatin accessibility and enhancer programs; conditional deletion in committed cDC1s drives transcriptional reprogramming toward cDC2-like states ^6–9^. IRF8 also governs type I interferon production and antigen presentation in pDCs, and IRF8-containing transcriptional circuits drive specialized anti-tumor DC subsets that promote durable T cell immunity ^10–12^. IRF8-dependent programs in tumor-associated macrophages and DCs likewise regulate antigen cross-presentation, CD8⁺ T cell priming, and anti-tumor immune memory, positioning IRF8 as a central regulator of myeloid cell-mediated tumor immunity ^13^.

IRF8 executes its regulatory functions through combinatorial interactions with lineage-defining transcription factors including PU.1, BATF3, SPIB, IRF1, IRF4, and AP-1/ETS family members ^14,15^. These partnerships enable context-dependent binding to distinct cis-regulatory motifs, canonical ISREs, ETS-IRF composite elements (EICEs), and AP1-IRF composite elements (AICEs) ^16–18^. In cDC1s, high IRF8 levels selectively engage AICE-containing enhancers to establish DC identity and antigen presentation programs ^7,19^. In microglia and pDCs, IRF8-dependent chromatin accessibility precedes transcriptional activation, demonstrating that IRF8 functions as an upstream chromatin organizer that licenses immune-responsive gene programs ^20^.

IRF8 is expressed throughout B cell development and participates in transcriptional networks with PU.1, IKAROS, E2A, EBF1, and PAX5 to govern B cell lineage specification and commitment ^21,22^. IRF8 directly regulates Sfpi1 and Ebf1, influencing early lineage differentiation ^21^. IRF8 also controls germinal center formation, marginal zone versus follicular fate decisions, immunoglobulin light chain rearrangement, and plasma cell differentiation ^23,24^. Cooperatively with PU.1, IRF8 represses BLIMP1-dependent plasma cell programs, and CRISPR-mediated IRF8 deletion promotes premature plasmablast formation and dysregulated plasma cell gene expression ^24^. Beyond normal B cell development, IRF8 dysregulation also shapes antigen presentation in malignant B cells^25^. IRF8 mutations in diffuse large B cell lymphoma disrupt CD74 and HLA-DM expression, impairing MHC class II-restricted antigen processing and remodeling the tumor immune microenvironment toward immune evasion^25^, underscoring that IRF8 is a B cell-intrinsic determinant of antigen-presenting competence even outside the myeloid compartment^25^. Despite these extensive studies, whether IRF8 governs functional activation programs of tumor-associated B cells remains unknown. B cells are now recognized as critical regulators of anti-tumor immunity ^26–29^. Tumor-infiltrating B cells and plasma cells are associated with improved responses to immune checkpoint blockade and favorable prognosis across multiple cancer types ^30–32^. Beyond antibody production, B cells function as professional APCs that activate CD4⁺ and CD8⁺ T cells through MHC-mediated antigen presentation, costimulatory molecule expression, and cytokine secretion ^33,34^. Notably, blocking plasma cell differentiation enhances antigen presentation by tumor-associated B cells and improves anti-tumor immunity, highlighting the critical importance of B cell differentiation state in regulating T cell activation within the TME ^28^.

Despite these advances, whether IRF8 establishes chromatin accessibility and antigen presentation programs in tumor-infiltrating B cells, the cis-regulatory motifs governing these states, and whether IRF8 acts cell-intrinsically rather than through myeloid functions remain unresolved. Here, using single-cell multiomics and B cell-specific deletion, we show IRF8 establishes a conserved chromatin program across myeloid cells and plasmablasts, governed by EICE rather than ISRE motifs in B cells, and required for antigen presentation and tumor growth control. This program is enriched in melanoma and breast cancer immunotherapy patient responders, establishing the IRF8-B cell axis as a clinically relevant regulator of anti-tumor immunity.

## RESULTS

### IRF8 deficiency accelerates tumor growth and reshapes the immune landscape

Loss of IRF8 impairs myeloid cell differentiation and promotes tumor growth ^1,35,36^. To elucidate the molecular mechanism underlying IRF8 function in anti-tumor immunity, we inoculated wild-type (WT) and *Irf8*-deficient (IRF8 KO) mice with B16-F10 tumor cells. Tumors developed in all IRF8 KO mice, whereas only 3 out of 5 WT mice formed measurable tumors (Fig. 1A & B). Tumor growth was significantly accelerated in IRF8 KO mice compared to WT mice (Fig. 1C & D). Within the CD45⁺ cell compartment, the frequencies of major lymphoid populations, including CD3⁺ T cells, CD19⁺ B cells, NK cells, and innate lymphoid cells (ILCs; CD90.2⁺), were all significantly decreased in IRF8 KO mice (Fig. 1E & F). These findings validate a broad impairment in lymphoid cell infiltration within IRF8 KO tumors. Systemically, tumor-bearing IRF8 KO mice developed splenomegaly (Fig. S1A), with significantly increased spleen size and total splenocyte numbers compared to WT mice (Fig. S1B). Despite this expansion, the relative frequencies of CD3⁺ T cells, CD19⁺ B cells, NK1.1⁺ NK cells, and CD90.2⁺ ILCs were significantly reduced in the spleens of IRF8 KO mice (Fig. S1C-D). Together, these results demonstrate that IRF8 deficiency profoundly reshapes both local and systemic immune landscapes, resulting in impaired lymphoid representation and defective anti-tumor immunity, consistent with IRF8 function in regulating myeloid and lymphoid immune homeostasis under pathological conditions ^13,37,38^.

**Figure 1.**
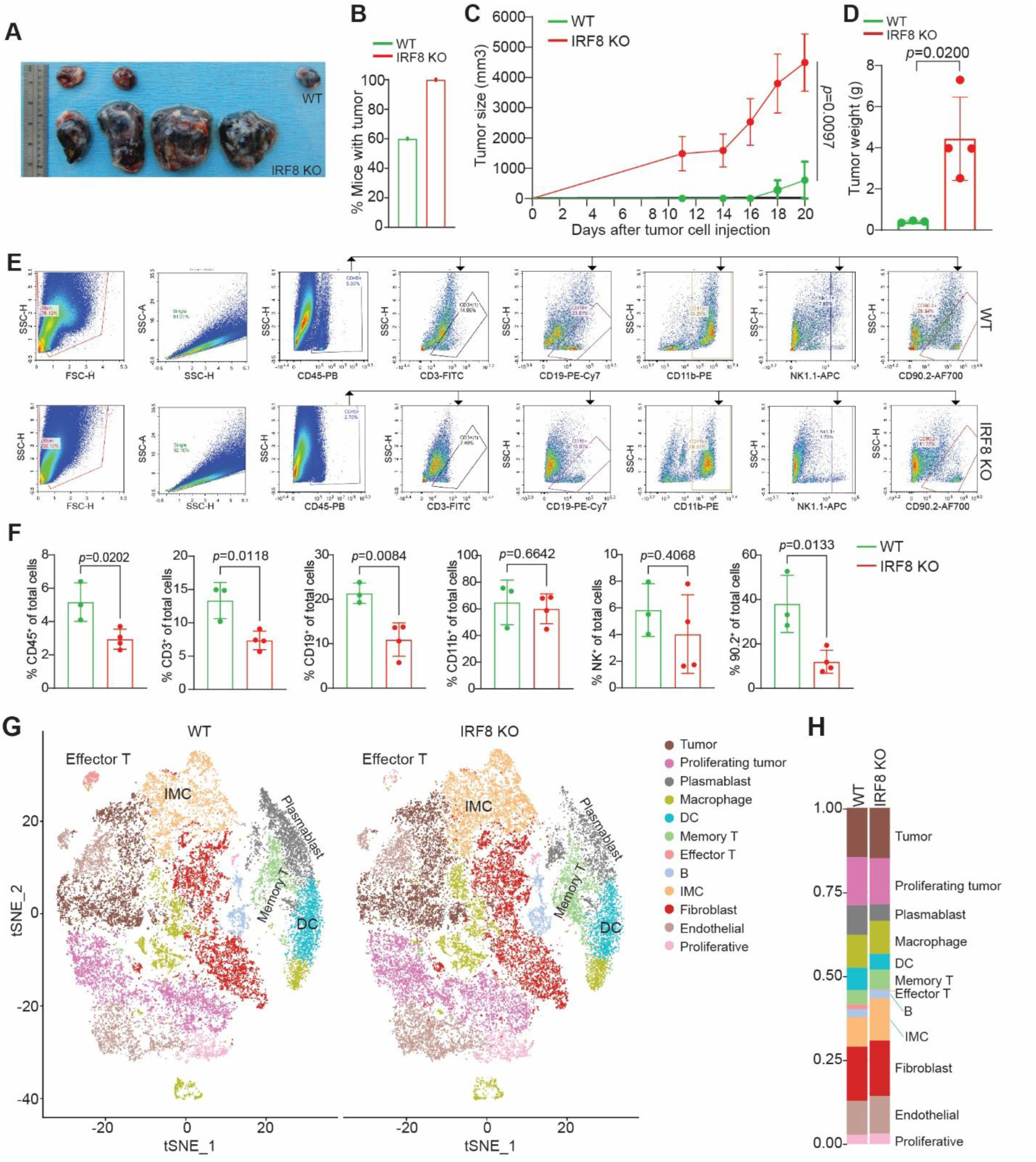
Single-cell transcriptomic profiling reveals IRF8-dependent remodeling of the tumor immune microenvironment. **A.** Images of B16-F10 tumors from WT and Irf8 KO mice. **B.** Tumor incidence in WT and IRF8 KO mice following subcutaneous B16-F10 implantation. **C.** Tumor growth kinetics in WT and IRF8 KO mice. **D.** Tumor weights at experimental endpoint. **E.** Representative flow cytometry gating strategy for tumor-infiltrating immune populations in WT and IRF8 KO B16-F10 tumors. **F.** Quantification of tumor-infiltrating immune cell populations shown in panel E. **G.** UMAP visualization of integrated scRNA-seq datasets showing cell populations identified in WT and IRF8 KO tumors. **H.** Bar plot quantification of relative cell composition in WT and IRF8 KO tumors.

### Single-cell transcriptomics reveals preserved lineage diversity but altered immune composition in the tumor microenvironment in IRF8 KO mice

The above findings prompted us to define immune cell heterogeneity and lineage distribution at single-cell resolution. Because IRF8 is a transcription factor, we isolated immune cells from tumors of WT and IRF8 KO mice and performed single-cell multiomics to couple transcription and chromatin remodeling in the same cell. Unsupervised clustering and UMAP visualization identified multiple immune populations, including effector T cells, memory T cells, dendritic cells (DCs), macrophages, myeloid-derived suppressor cells (IMCs), B cells, plasmablasts, NK cells, neutrophils, endothelial cells, and fibroblasts (Fig. 1G & H). Overall cellular identities were preserved between WT and IRF8 tumors, indicating that IRF8 loss does not eliminate major immune lineages but instead reshapes their relative abundance and perhaps functional states within the tumor microenvironment. Comparative analysis revealed marked compositional changes in IRF8 KO tumors. Most notably, IMCs were substantially expanded in IRF8 KO tumors, consistent with the established role of IRF8 in restraining immature myeloid cell accumulation^35,36^. In contrast, DCs, plasmablasts, and effector T cells were markedly reduced in IRF8 KO tumors (Fig. 1G & H). Interestingly, although total T cell populations were decreased, IRF8 loss resulted in a relative enrichment of memory-like T cells accompanied by a reduction in cytotoxic effector T cells, suggesting impaired effector differentiation rather than T cell exclusion.

### A conserved IRF8-dependent antigen presentation signature spans myeloid and B cell lineages

Our above analysis revealed substantial alterations in plasmablasts, DC, and IMC in IRF8 KO tumors compared to WT controls. While IRF8 is well established as a master regulator of myeloid lineage differentiation and antigen-presenting cell function ^16,39–42^, its role in B cell functional programming within tumors remains poorly understood. To determine whether IRF8 regulates B cell function in a manner analogous to its canonical role in myeloid cells, we used DCs and IMCs as reference antigen-presenting populations ^43,44^ and performed comparative differential gene expression analyses across DCs, IMCs, and plasmablasts isolated from WT and IRF8 KO tumors (Fig. 2A-F). Across all three immune populations, differential expression analyses identified a strikingly conserved IRF8-dependent transcriptional program centered on antigen processing and presentation pathways (Fig. S2). Genes involved in MHC class I and antigen presentation machinery, including Cd74, H2-D1, H2-K1, B2m, Tap1, and Psmb8, were consistently reduced in IRF8 KO cells (Fig. 2A-I). In parallel, interferon-response regulators such as Stat1, Bst2, and Iigp1 were coordinately suppressed, indicating impaired IFN-associated immune activation programs in the absence of IRF8 (Fig. 2G-I). These shared transcriptional alterations across DCs, IMCs, and plasmablasts, together with overlapping differentially expressed genes (Fig. 2J) suggest that IRF8 functions as a conserved regulator of antigen presentation across both myeloid and B lineage compartments.

**Figure 2.**
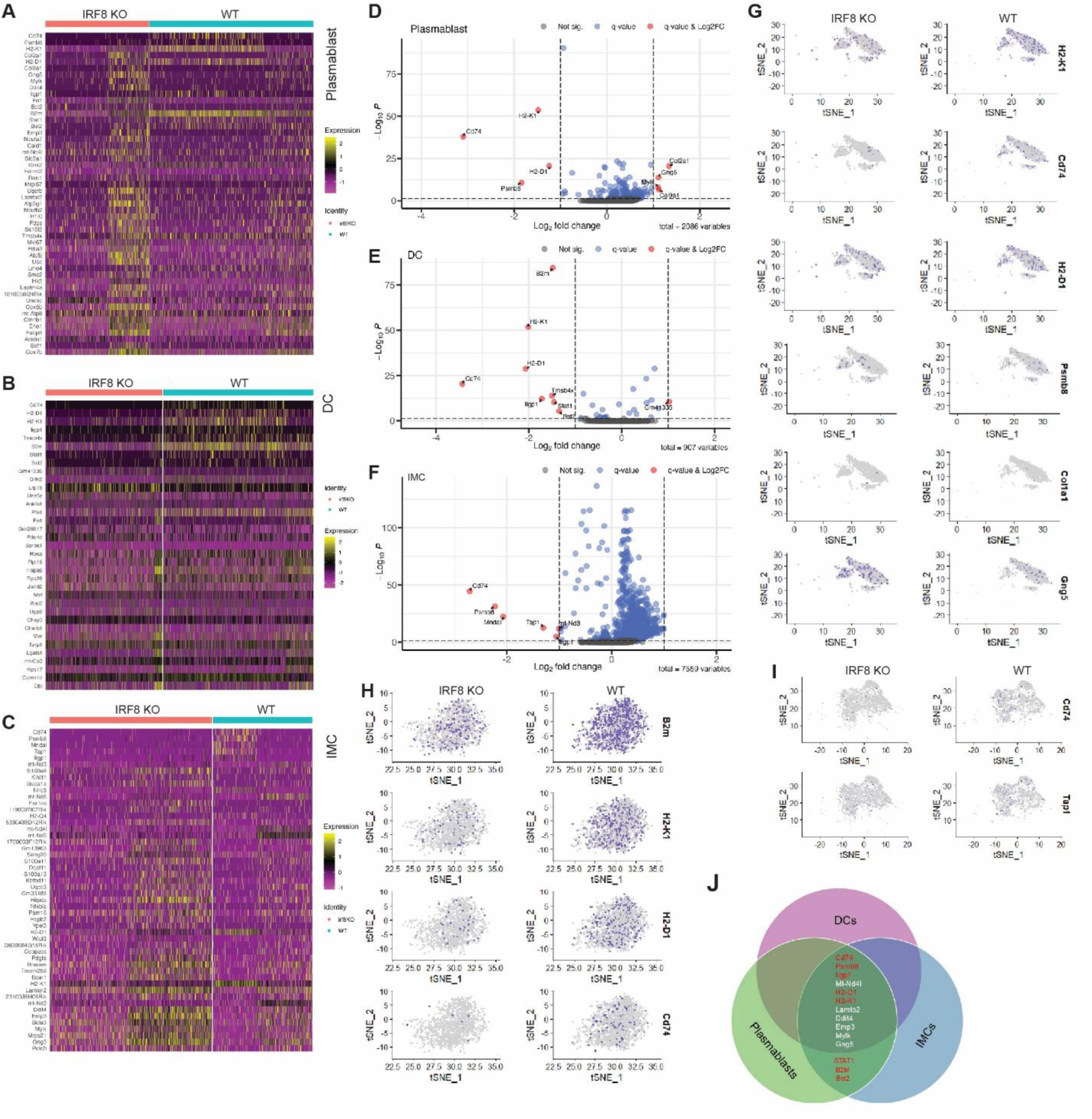
IRF8 regulates a shared antigen presentation transcriptional program across plasmablasts, DCs, and IMCs. **A-C.** Heatmaps showing differentially expressed genes between WT and IRF8 KO cells in plasmablasts (A), DCs (B), and IMCs (C). Genes associated with antigen processing and presentation are highlighted with red boxes. **D-F**. Volcano plots of differential gene expression analyses comparing WT and IRF8 KO plasmablasts (D), DCs (E), and IMCs (F). Significantly upregulated and downregulated genes are highlighted. **G-I**. UMAP feature plots showing expression of representative antigen presentation and interferon-response genes in WT and IRF8 KO plasmablasts (G), DCs (H), and IMCs, (I). **J**. Venn diagram showing overlap of differentially expressed genes among plasmablasts, DCs, and IMCs.

In addition to antigen presentation defects, IRF8 KO cells also exhibited transcriptional signatures consistent with metabolic and stress adaptation. Differentially expressed mitochondrial and oxidative phosphorylation genes, including mt-Nd, Cox, Atp5, and Uqcrb, suggested metabolic rewiring across myeloid and plasmablast populations, whereas stress- and hypoxia-associated genes such as Ddit4, Hilpda, and Slc2a1 were increased in Irf8-deficient cells (Fig. 2D-F). IMCs additionally displayed enrichment of proliferation-associated genes including Mki67, Ect2, Smc2, and Rrm1, consistent with accumulation of immature myeloid populations in IRF8 KO tumors.

### Distinct IRF8-dependent motif architectures define chromatin accessibility programs across tumor-infiltrating myeloid cells and plasmablasts

A panel of 38 IRF8-like motifs, spanning IRF, ISRE, EICE/AICE composite, PU.1/ETS families, was curated and interrogated in tumor-infiltrating DCs, IMCs, and plasmablasts from WT and IRF8 KO mice (Table S1). Motif enrichment analysis of IRF8-associated chromatin accessibility peaks across these three cell types reveals both shared and lineage-specific regulatory architectures that are differentially sensitive to IRF8 loss (Fig. S3). In WT DCs, standalone ISRE and IRF motifs are depleted at accessible IRF8-like peaks, while ETS-family and composite ETS:RUNX motifs predominate, suggesting that DCs preferentially exploit composite regulatory elements rather than canonical ISRE half-sites (Table S2). Notably, AP-1/bZIP motifs are also strongly depleted in WT DC accessible peaks, indicating that IRF8 operates in DCs through an AP-1-independent, ETS-composite mode. In WT IMCs, the chromatin accessible at IRF8-like loci is saturated with canonical IRF/ISRE motifs, including ISRE, T1ISRE, IRF1, IRF2, bZIP:IRF (AICE), and JASPAR IRF8 matrices, alongside a strikingly strong AP-1/bZIP co-enrichment signature, reflecting an active, broad-spectrum IRF8-centered transcriptional program tightly integrated with AP-1 coregulators. Notably, WT plasmablasts occupy a distinct regulatory niche: the dominant motif enrichment signature involves ETS/PU.1-family elements together with specific EICE composite motifs (PU.1-IRF and MA1484.1), with AP-1/bZIP accessibility strongly depleted (Table S4), indicating that IRF8 operates primarily in a PU.1-cooperative, EICE-dependent mode in B lineage cells rather than through the standalone ISRE program dominant in myeloid cells ^20^. Consistent with this, ISRE enrichment does not reach significance in WT plasmablasts, further distinguishing the B cell IRF8 regulatory program from the canonical myeloid ISRE-dominated architecture.

Upon IRF8 deletion, the motif landscapes shift in cell population-specific ways. In DCs, the residual IRF/ISRE accessibility is further eroded and canonical IRF8 (MA0652.1)(Fig. S3) is significantly lost, while ETS-family enrichment is maintained or amplified (Table S5). In IMCs, the overall IRF motif enrichment is broadly preserved despite IRF8 loss (Table S6), a finding consistent with the massive expansion of IMCs in the IRF8 KO mice ^13,35^. Most strikingly, in IRF8 KO plasmablasts, the IRF component of the accessible chromatin landscape is essentially abolished: ISRE, T1ISRE, IRF1, IRF2, IRF4, PU.1-IRF, and bZIP:IRF motifs all lose significance, while the ETS scaffold persists (Table S7). Importantly, MA1484.1, an EICE composite element containing both ETS and IRF half-sites, remains significantly enriched in IRF8 KO plasmablasts at levels comparable to WT. This persistence likely reflects continued occupancy of the ETS half-site by PU.1 in the absence of IRF8, rather than intact EICE binding, since the canonical PU.1-IRF composite motif is lost in the KO. Taken together, these findings indicate that IRF8 loss selectively ablates the IRF component of EICE-dependent chromatin accessibility in plasmablasts, while PU.1-anchored ETS occupancy persists independently of IRF8. These data reveal that IRF8 is the critical determinant of the IRF/EICE regulatory program at accessible chromatin in plasmablasts, that it contributes to, though is not solely responsible for, IRF/ISRE chromatin accessibility in DCs, and that in IMCs, the IRF8-like chromatin program reflects a population-level phenomenon partially maintained by the expanded immature myeloid compartment in the absence of IRF8.

### IRF8 establishes lineage-spanning chromatin accessibility programs in myeloid and B cells

To map chromatin accessibility landscapes in WT and IRF8 KO immune cells, we next integrated single-cell ATAC-seq and RNA-seq data from DCs, IMCs, and plasmablasts. Dimensionality reduction and UMAP embedding resolved the major immune populations and demonstrated genotype-dependent chromatin remodeling within each lineage (Fig. S4A). Genome-wide annotation of accessible peaks containing IRF8-like motifs revealed that the majority localized to intronic and distal intergenic regions rather than promoter-proximal elements, consistent with enhancer-mediated regulation and conserved across all three lineages (Fig. S4B). DA analysis identified significantly remodeled peaks per lineage (Fig. S4C-K). In DCs, 1,024 peaks showed increased and 2,270 peaks showed decreased accessibility in IRF8 KO relative to WT cells (Fig. S4C-E). Comparable changes were observed in IMCs (958 increased, 1,270 decreased; Fig. S4F-H) and plasmablasts (1,097 increased, 1,825 decreased; Fig. S4I-K). Peaks harboring IRF8-like motifs were significantly enriched among differentially accessible loci in all three cell types (Fig. S4D, G, J). Notably, plasmablasts displayed extensive IRF8-dependent chromatin remodeling comparable in scale to that seen in canonical myeloid antigen-presenting cells (Fig S4I-K), supporting a direct role for IRF8 in B cell functional programming within the tumor microenvironment.

### IRF8 regulates conserved enhancer and motif architectures across myeloid and B lineage cells

Genome-wide annotation of accessible IRF8-like peaks revealed that a large portion of IRF8-associated chromatin regions localized to intronic and distal intergenic regions rather than promoter-proximal elements (Fig. 3A & B), consistent with enhancer-mediated transcriptional regulation ^45,46^. This enhancer-biased genomic distribution was observed across DCs, IMCs, and plasmablasts in both WT and IRF8 KO tumors, indicating that IRF8 regulates both distal cis-regulatory elements and core promoters regardless of lineage context. Differential accessibility analyses revealed widespread chromatin remodeling in IRF8 KO immune cells, with more accessibility peaks gained than lost in IRF8 KO DCs, IMCs, and plasmablasts relative to WT controls (Fig. 3C & D, Fig. S3A-F). Analysis of DA peak distribution relative to TSSs demonstrated enrichment of accessibility changes within enhancer-associated distal genomic regions across all three populations (Fig. 3E), further supporting the concept that IRF8 regulates both enhancer and promoter-restricted accessibility landscapes. Density plots of inter-peak distances showed comparable chromatin compaction patterns in WT and IRF8 KO for all three cell types (Fig. 3F), suggesting that the gross spatial organization of IRF8-associated regulatory elements is preserved even as individual sites gain or lose accessibility. MA and volcano plots of DA results, with IRF8-like motif-containing peaks highlighted as gained or lost, confirmed substantial bidirectional remodeling (Fig. 3G). Notably, plasmablasts exhibited IRF8-dependent chromatin remodeling of comparable magnitude to that observed in canonical antigen-presenting myeloid populations, providing additional evidence that IRF8 directly regulates B cell transcriptional programming within the tumor microenvironment.

**Figure 3.**
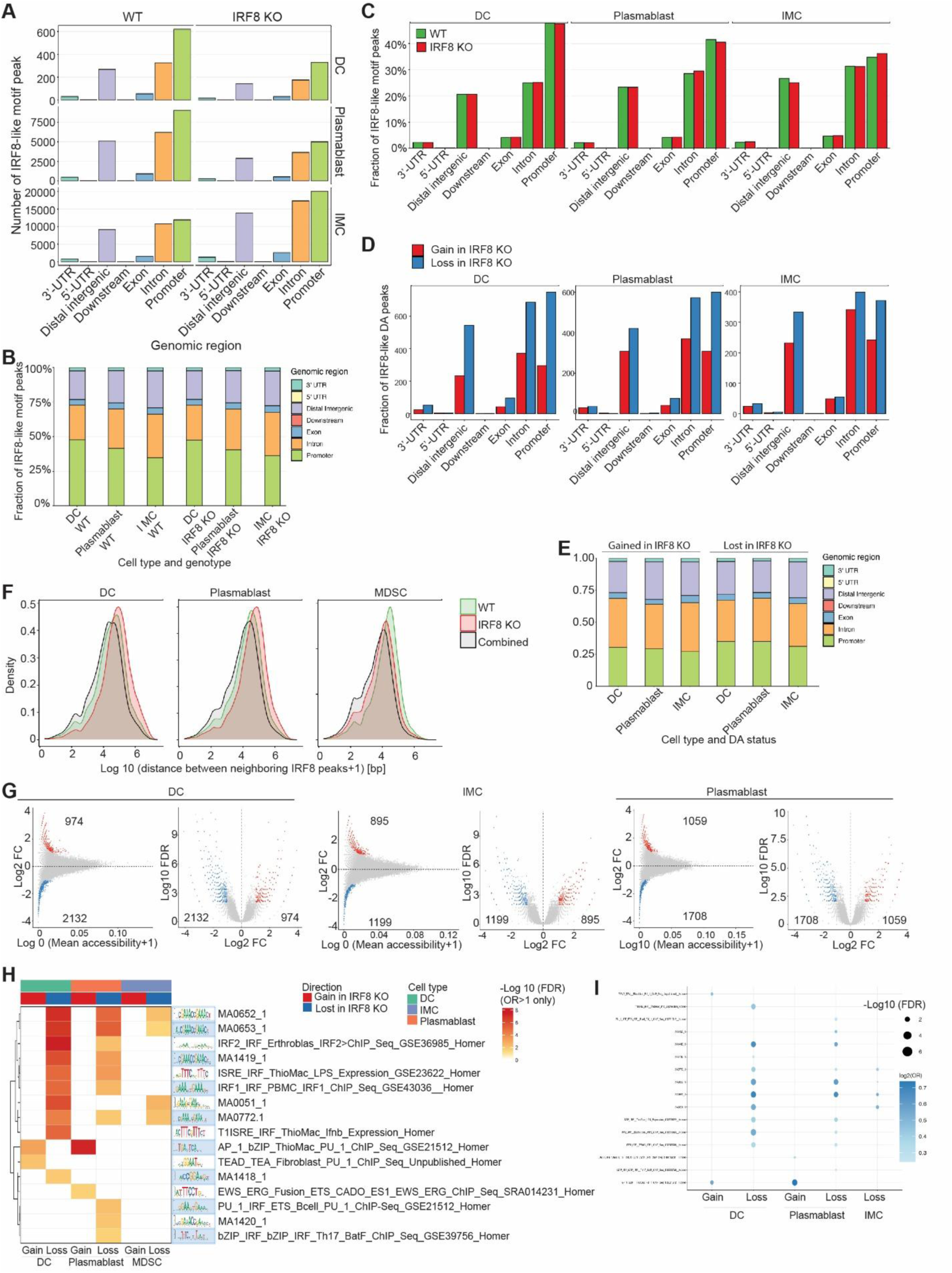
IRF8-mediated chromatin accessibility regulates antigen presentation pathways across myeloid and plasmablast cells. A. Genomic annotation of accessible IRF8-like peaks in DCs, IMCs, and plasmablasts from WT and IRF8 KO tumors. Peaks harboring at least one match to any of the IRF8-like PWMs were annotated with ChIPseeker (TxDb.Mmusculus.UCSC.mm10.knownGene; TSS window ±3 kb). Bars show the absolute number of IRF8-like peaks falling within promoter, 5′UTR, exon, intron, 3′UTR, downstream, and distal intergenic categories per cell type and genotype. B. Stacked bar chart of the fractional genomic distribution of IRF8-like peaks across all cell type and genotype groups. C. Bar graphs quantifying the total number of DA chromatin peaks gained (KO>WT, log₂FC> 1, FDR<0.01) and lost (WT>KO, log₂FC<−1, FDR< 0.01) in IRF8 KO cells relative to WT controls. DA was computed using a binomial test on binarized peak accessibility matrices within per-cell-type ArchR subprojects. D. Grouped Bar plots show the number of IRF8-like motif-containing DA peaks annotated to each genomic region category in DC, Plasmablast, and IMC cells. Red bars indicate peaks with significantly increased accessibility in IRF8 KO relative to WT (gained in KO; log₂FC>1, FDR<0.01); blue bars indicate peaks with significantly reduced accessibility (lost in KO; log₂FC<-1, FDR<0.01). Genomic annotations were assigned using ChIPseeker with a ±3 kb TSS window. E. Stacked bar chart of the genomic annotation distribution of significantly DA IRF8-like peaks across all three cell types. F. Density plots of log₁₀-transformed distances between consecutive neighboring IRF8-like peaks in WT, IRF8 KO, and combined cells for DCs, plasmablasts, and IMCs. G. MA plots and volcano plots of DA analyses for DCs, IMCs, and plasmablasts. Gray points represent all tested peaks. Colored points indicate IRF8-like peaks passing significance thresholds (FDR<0.01, log₂FC≥1: red=gained accessibility in KO (log₂FC >1); blue=lost accessibility in KO (log₂F <-1). Peak counts for each category are indicated within each panel. H. Heatmap of IRF8-like motif enrichment within DA peak sets across cell types and DA direction. Motif annotations are shown alongside published ChIP-seq identifiers where available. I. Dot plot summarizing IRF8-like motif enrichment patterns associated with gained and lost IRF8-dependent DA peaks in DCs, plasmablasts, and IMCs.

Having established that each immune population deploys a distinct IRF8 regulatory element with ISRE-dominant in IMCs, ETS-composite in DCs, and EICE-dependent in plasmablasts, we next asked whether these lineage-specific programs converge on a shared structural feature at the level of motif architecture. A heatmap of enrichment significance across IRF8-like motif variants and all cell-type and DA-direction combinations (Fig. 3H, Fig. S5) and a complementary dot plot of enrichment statistics (Fig. 3I) revealed that ETS/PU.1-family elements represent the most broadly shared component of IRF8-dependent chromatin accessibility across all three populations, with the EICE being the only motif class achieving significant enrichment in DCs, IMCs, and plasmablasts alike. Together, these data indicate that a PU.1-anchored ETS scaffold constitutes the conserved structural core of IRF8-dependent regulatory elements across immune lineages, onto which lineage-specific IRF co-factor interactions, ISRE-class in myeloid cells, EICE-class in B lineage cells, are superimposed to generate population-adapted chromatin accessibility program.

### Integrated ATAC-RNA analyses identify coordinated IRF8-dependent transcriptional programs in antigen-presenting immune cells

To determine whether IRF8-dependent chromatin accessibility remodeling correlates with transcriptional changes, we performed integrated ATAC-RNA analyses linking IRF8-like DA peaks with their nearest DE genes across DCs, plasmablasts, and IMCs. IRF8-like DA peaks were first identified by binomial differential accessibility testing (FDR<0.01, log₂FC≥1) within each cell type, then cross-referenced against RNA DE results (Wilcoxon FD ≤0.01, log₂FC ≥0.58) to retain only peak-gene pairs in which the nearest gene was independently regulated at the transcriptional level. This dual-stringency filtering yielded 209 peak-gene pairs in DC, 171 in plasmablasts, and 163 in IMC (Fig. 4A-C). Heatmap analysis centered on IRF8-like DA peaks (±1 kb windows, 10-bp bins) demonstrated coordinated genotype-dependent changes in chromatin accessibility and gene expression across all three immune populations (Fig. 4A-C). Within each cell type, DA peaks were partitioned into up and down clusters; rows within each cluster were ordered by the magnitude of the KO-WT RNA expression difference of the nearest DE gene, enabling visual alignment of chromatin and transcriptional changes. Regions exhibiting reduced chromatin accessibility in IRF8 KO cells, the predominant class across all three populations, were associated with concomitant decreases in expression of nearby genes, consistent with IRF8-dependent enhancer accessibility contributing to the maintenance of transcriptional activation programs across myeloid and B lineage compartments. The aggregated enrichment profiles above the heatmaps confirmed a clear shift from symmetric accessibility peaks in WT to attenuated or absent signals in IRF8 KO cells at down-cluster loci, while up-cluster loci showed the reciprocal pattern of newly acquired accessibility in the IRF8 KO context (Fig. 4A-C).

**Figure 4.**
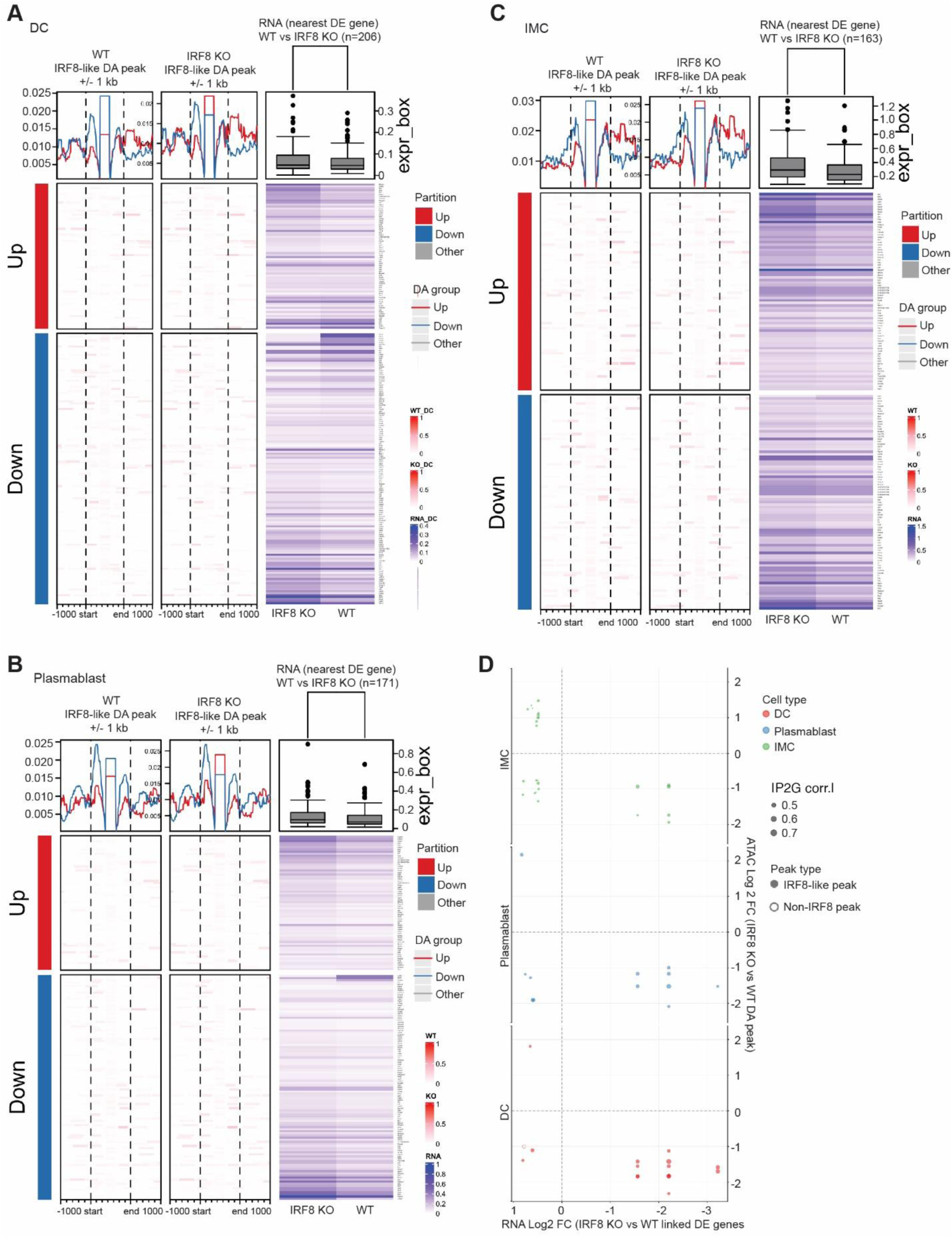
Integrated ATAC-RNA analyses reveal coordinated IRF8-dependent chromatin accessibility and transcriptional regulation across myeloid and B lineage cells. **A-C**. Heatmaps showing integrated differential chromatin accessibility and transcriptional analyses centered on IRF8-like DA peaks and their nearest DE genes in DCs (A; n=209 peak-gene pairs), plasmablasts (B; n=171 peak-gene pairs), and IMCs (C; n=163 peak-gene pairs) from WT and IRF8 KO tumors. DA peaks were defined by binomial testing. Peaks are split into "Up" (gained accessibility in IRF8 KO; red) and "Down" (lost accessibility in IRF8 KO; blue) clusters; within each cluster, rows are ordered by the magnitude of the KO-WT RNA expression difference of the nearest DE gene. Left: DA group partition annotation (red=Up, blue=Down). Middle: ATAC accessibility enrichment heatmaps showing WT and IRF8 KO signal across ±1 kb windows centered on IRF8-like DA peak summits (bin size=10 bp), with aggregate enrichment profiles displayed above. Right: RNA expression heatmaps of the nearest DE gene associated with each IRF8-like DA peak in WT and IRF8 KO cells; boxplots above show the distribution of average log-normalized expression values. **D**. Integrated ATAC-RNA scatter plot showing relationships between chromatin accessibility changes and transcriptional changes across DCs, plasmablasts, and IMCs. Each point represents a Peak2Gene-linked IRF8-like DA peak-DE gene pair; bubble size reflects the Peak2Gene correlation coefficient. IRF8-like DA peaks are shown as filled circles; non-IRF8-associated DA peaks are shown as open circles. Positive correlations between ATAC and RNA fold changes indicate coordinated regulation; the tighter clustering of IRF8-like peaks along the diagonal relative to non-IRF8 peaks reflects the specificity of IRF8-centered regulatory elements in driving transcriptional output.

These chromatin accessibility alterations were tightly linked to reduced expression of nearby genes involved in antigen processing, interferon signaling, and immune activation pathways, including members of the MHC class I presentation machinery (e.g., H2-D1, B2m, Tap1). The RNA expression heatmaps (Fig. 4A-C, right panels) and their accompanying boxplots confirm the directional concordance between ATAC and RNA changes. The top-ranked rows within the down cluster, sorted by largest KO-WT RNA difference, consistently displayed both the most prominent loss of chromatin accessibility and the greatest transcriptional suppression.

While plasmablasts share the functional output of IRF8-dependent chromatin remodeling with DCs and IMCs, including coordinated loss of accessibility and transcriptional suppression at antigen presentation-associated loci, the underlying regulatory architecture is distinct. plasmablasts deploy an EICE-dependent regulatory program rather than the ISRE-dominant architecture of IMCs or the ETS-composite program of DCs (Table S2-7, Fig. 3). The convergence on a shared functional consequence, suppression of antigen processing and MHC class I presentation machinery, through population-adapted chromatin mechanisms provides strong evidence that IRF8 licenses antigen-presenting competence across immune lineages through lineage-specific epigenetic programs rather than a single uniform regulatory circuit (Fig. 3, Fig. S4).

To globally assess the relationship between chromatin accessibility and gene expression, we integrated ATAC accessibility fold changes with RNA expression fold changes across all three cell types using Peak2Gene-linked pairs (Peak2Gene correlation≥0.45, FDR≤0.05; Fig. 4D). Scatter plot analyses demonstrated a positive correlation between reduced chromatin accessibility and decreased transcriptional output in IRF8 KO cells, consistent across DCs, plasmablasts, and IMCs (Fig. 4D). IRF8-like peaks clustered more tightly along the diagonal than non-IRF8-associated DA regions, and the Peak2Gene correlation coefficients were systematically higher for IRF8-like loci (Fig. 4D). This indicates that IRF8-centered regulatory elements are primary contributors to immune transcriptional programs in the tumor microenvironment rather than incidental bystanders to broader chromatin remodeling, and that the strength of the ATAC-RNA coupling is itself a distinguishing feature of IRF8-bound regulatory elements.

### IRF8-associated chromatin accessibility programs directly regulate antigen presentation gene expression across myeloid and plasmablast populations

To define the functional consequences of IRF8-dependent chromatin accessibility remodeling, we integrated DA peaks with Peak2Gene linkages and transcriptional analyses across DCs, plasmablasts, and IMCs. GO enrichment analyses of DE genes associated with IRF8-like DA peaks revealed significant enrichment of antigen processing and presentation, interferon signaling, immune activation, and inflammatory response pathways across all three immune populations (Fig. 5A). In DCs and plasmablasts, antigen processing and presentation emerged as the top-ranked enriched biological pathway, consistent with the categorical loss of IRF-family motif accessibility observed in these populations upon IRF8 deletion. In IMCs, antigen presentation-associated GO terms were also enriched among IRF8-linked DE genes, though with comparatively smaller gene counts than in DCs or plasmablasts, reflecting partial rather than complete attenuation of the IRF program in this population.

**Figure 5.**
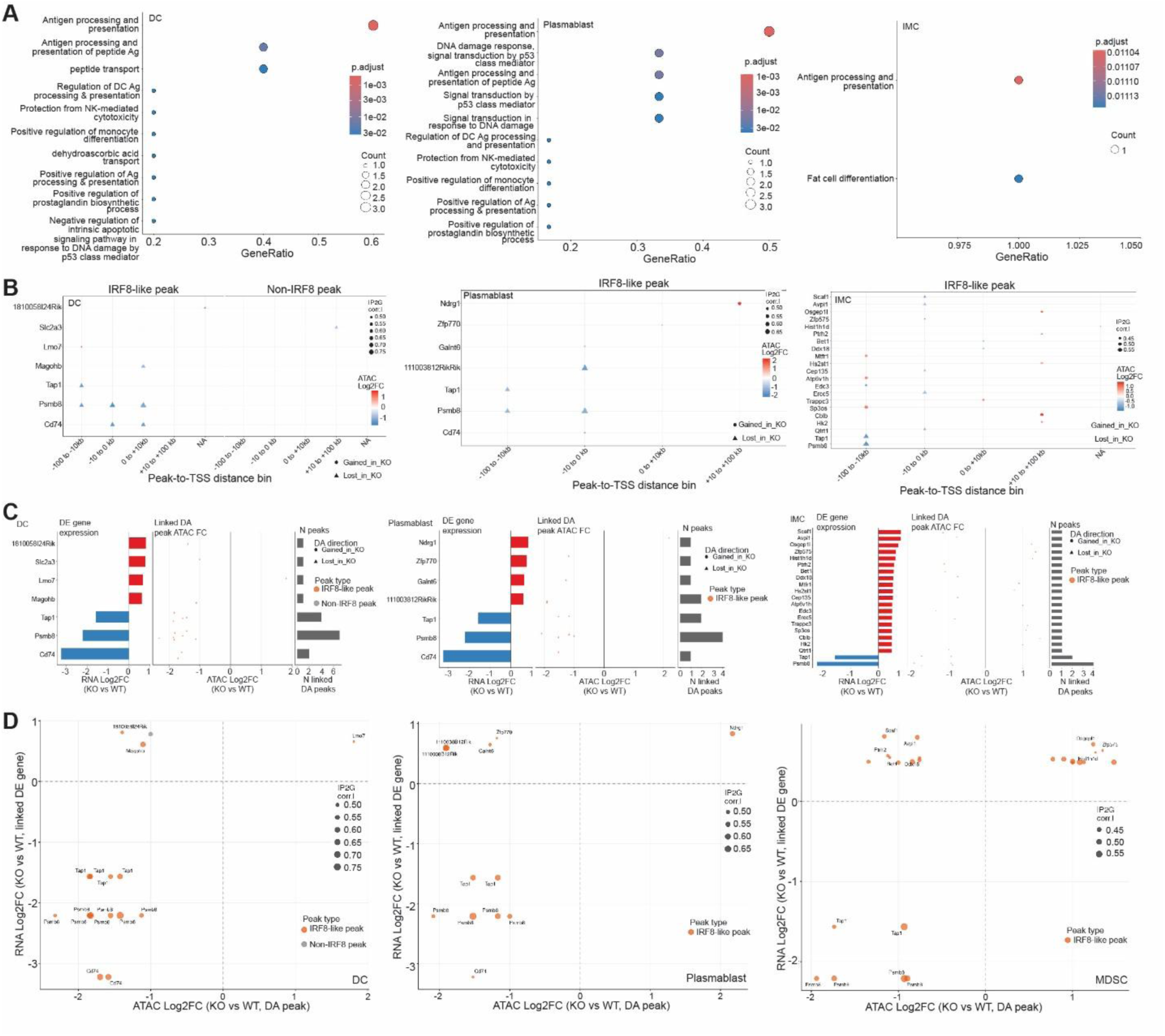
IRF8-dependent chromatin accessibility regulates antigen presentation gene expression programs across myeloid and B lineage cells. A. Dot plots of Gene Ontology (GO) Biological Process enrichment analyses of DE genes linked to IRF8-like DA peaks via Peak2Gene associations in DCs (left), plasmablasts (middle), and IMCs (right). The top 10 enriched terms per cell type are shown, ranked by gene count. B. Dot-matrix plots showing the distribution of IRF8-like DA peaks (left panels, orange) and non-IRF8 DA peaks (right panels, gray) linked to DE genes via Peak2Gene associations in DCs, plasmablasts, and IMCs. The y-axis represents DE genes ranked by RNA log₂ fold change (IRF8 KO vs WT), and the x-axis indicates peak-to-TSS distance bins (<-100 kb, - 100 to -10 kb, -10 kb to 0, 0 to +10 kb, +10 to +100 kb, >+100 kb). C. Integrated peak-to-gene annotation plots of RNA DE genes linked to DA chromatin peaks in DCs, plasmablasts, and IMCs, showing the top DE genes ranked by RNA log₂FC. Left bar panels: RNA expression log₂FC (IRF8 KO v WT) for each DE gene; red bars indicate upregulation and blue bars indicate downregulation in IRF8 KO. Middle strip panels: ATAC accessibility log₂FC of all DA peaks linked to each gene via Peak2Gene associations; IRF8-like peaks are shown in orange and non-IRF8-associated peaks in gray. Right bar panels: total number of DA peaks (IRF8-like and non-IRF8) linked to each DE gene, indicating the regulatory complexity at individual loci. Genes associated with antigen presentation and MHC class I machinery are highlighted across all three cell types. D. Scatter plots comparing ATAC differential accessibility log₂FC (x-axis; IRF8 KO vs WT, DA peak) and RNA differential expression log₂FC (y-axis; IRF8 KO vs WT, linked DE gene) for all Peak2Gene-linked DA peak-DE gene pairs in DCs (left), plasmablasts (middle), and IMCs (right). IRF8-like peaks are highlighted in orange; non-IRF8 DA peaks are shown in gray. Bubble size reflects the Peak2Gene correlation coefficient.

To characterize the spatial relationship between IRF8-dependent regulatory elements and their transcriptional targets, we analyzed peak-to-TSS distance distributions stratified by DE gene RNA level. Dot-matrix analyses demonstrated that IRF8-like DA peak-gene pairs cluster preferentially near the most highly differentially expressed immune regulatory genes (Fig. 5B, left panel), particularly within proximal distance bins. By contrast, non-IRF8 DA peaks distributed more broadly across DE genes spanning a wider range of fold-change magnitudes (Fig. 5B, right panels), with no preferential enrichment at the most strongly regulated genes. This spatial selectivity indicates that IRF8-centered cis-regulatory elements are specifically associated with the genes most severely affected by IRF8 loss, rather than contributing uniformly across the full DE gene landscape.

Integrated peak-to-gene annotation plots revealed coordinated relationships between chromatin accessibility and transcriptional output at individual gene loci (Fig. 5C). Multiple antigen presentation and interferon-response genes exhibited simultaneous reductions in RNA expression together with loss of accessibility at linked IRF8-like peaks in IRF8 KO cells, while non-IRF8-associated linked peaks showed comparatively modest or inconsistent accessibility changes at the same loci (Fig. 5C). This pattern was observed equivalently in plasmablasts, DCs, and IMCs, providing direct evidence that IRF8 regulates B cell antigen presentation programs through enhancer-associated chromatin accessibility. The number of IRF8-like DA peaks linked per gene further indicates that several antigen presentation loci are regulated by multiple convergent IRF8-dependent enhancer elements, suggesting that IRF8 coordinates regulatory hub architecture rather than single-enhancer control at these immunostimulatory genes.

Scatter plot analyses integrating chromatin differential accessibility with RNA differential expression confirmed a positive correlation between ATAC and RNA fold changes across DCs, plasmablasts, and IMCs (Fig. 5D). IRF8-like peaks displayed substantially tighter ATAC-RNA coupling compared to non-IRF8 peaks, indicating that IRF8-associated regulatory regions are primary determinants of immune gene expression programs in tumors. Together, these findings demonstrate that IRF8-dependent chromatin accessibility programs directly drive antigen presentation and immune activation gene expression across both myeloid and B lineage compartments, and that IRF8-centered enhancer elements represent the dominant functional drivers of this coordinated transcriptional regulation.

### B cell intrinsic IRF8 function sustains T cell activation and anti-tumor immunity in vivo

The above findings indicate that IRF8 intrinsically programs B cell antigen-presenting function within the tumor microenvironment, raising the question of whether tumor-associated B cells can directly activate T cells. To test this, we evaluated the ability of B cells to support T cell activation ex vivo. Splenic B cells and T cells isolated from tumor-bearing mice were co-cultured ex vivo. Flow cytometric analysis demonstrated that B cells induced T cell proliferation in a B cell:T cell ratio-dependent manner (Fig. 6A), directly supporting the capacity of tumor-bearing host B cells to function as antigen-presenting and T cell-activating populations.

**Figure 6.**
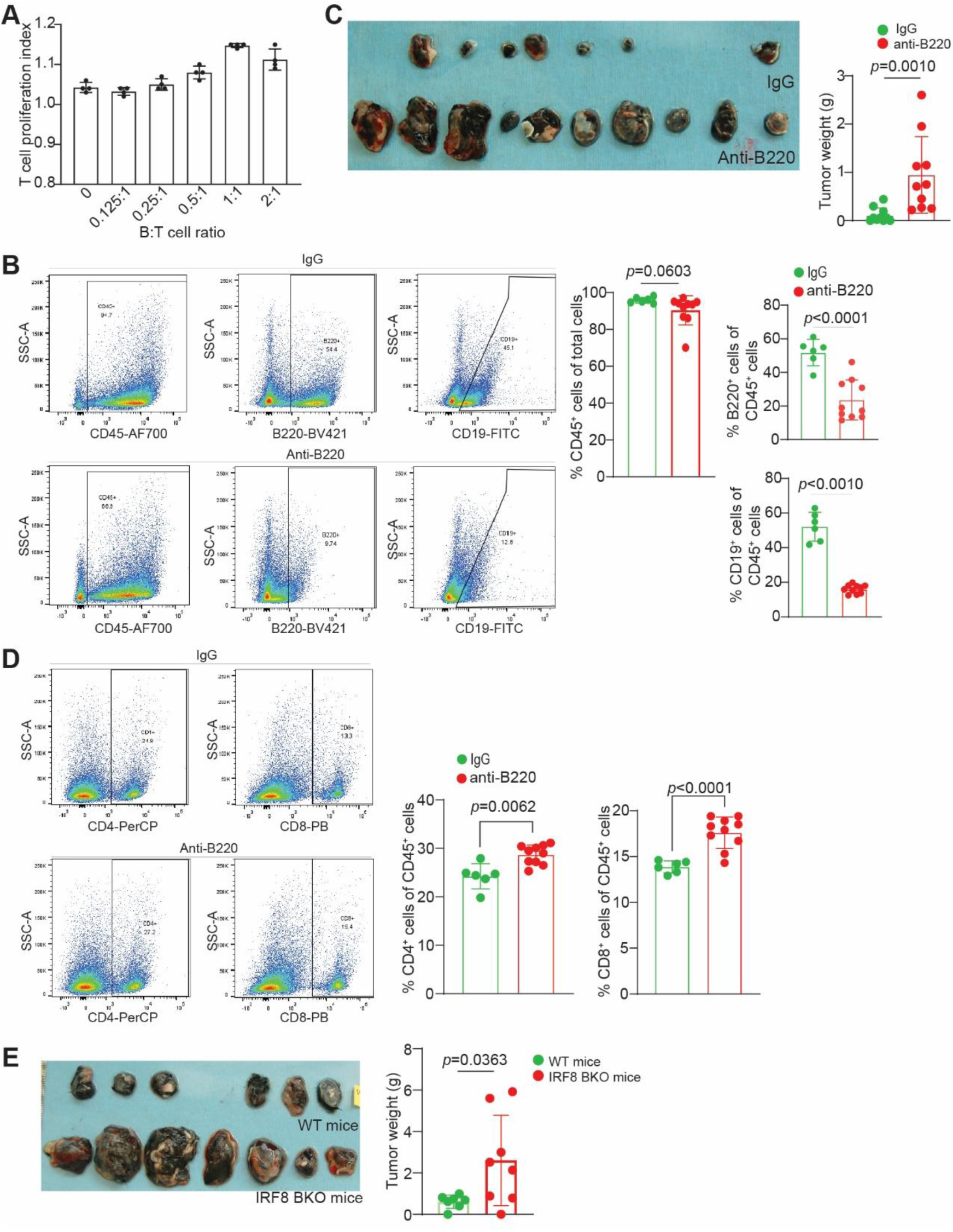
*Irf8* deletion in B cells impairs anti-tumor immunity and promotes tumor growth. **A.** Splenic B cells and T cells were isolated from tumor-bearing mice. T cells were labeled with CFSE and co-cultured with B cells at the indicated ratios for 3 days. T cell proliferation was analyzed by flow cytometry following gating on CD3⁺ T cells. **B.** B16 -F10 tumor cells were injected subcutaneously into mice followed by treatment with control IgG or B220-neutralizing monoclonal antibody every 3 days for a total of four treatments. Flow cytometry analysis of splenocytes showing depletion efficiency based on B220 and CD19 expression. **C.** Tumors from IgG- and B220-treated mice (left) and quantification of tumor weights at endpoint (right). **D.** Flow cytometry analysis of CD4^+^ and CD8^+^ T cells in spleens of IgG- and anti-B220-treated mice. Shown are gating (left) and quantification (right) **E.** B16 -F10 tumor cells were injected subcutaneously into WT mice (n=7) and mice with IRF8 deletion only in B cells (IRF8 BKO, n=7), followed by treatment with control IgG or B220-neutralizing monoclonal antibody every 3 days for a total of four treatments. Shown are tumor images at the endpoint (left) and tumor weight quantification.

To determine whether B cells contribute to tumor control in vivo, we performed antibody-mediated B cell depletion using an anti-B220 monoclonal antibody in tumor-bearing mice. In the B16-F10 melanoma model, mice were treated with anti-B220 antibody or control IgG following tumor implantation. Flow cytometric analyses confirmed effective depletion of splenic B220⁺ and CD19⁺ B cells in anti-B220-treated mice relative to controls (Fig. 6B). B cell depletion significantly increased tumor weight compared to IgG-treated controls (Fig. 6C). Consistent with the ex vivo evidence that B cells directly support T cell activation (Fig. 6A), B cell depletion significantly reduced tumor-infiltrating T cells in tumor-bearing mice (Fig. 6D), suggesting that the pro-tumorigenic effect of B cell depletion is mediated, at least in part, through impaired intratumoral T cell priming. To determine whether this phenotype was tumor model-dependent, we evaluated the effects of B cell depletion in the CT26 colon tumor model. Similar to the B16-F10 melanoma model, anti-B220 treatment significantly enhanced CT26 tumor growth and increased tumor mass relative to control-treated mice (Fig. S6A & B), establishing a non-redundant role for B cells in restraining tumor progression across distinct cancer types. To directly establish the B cell-intrinsic contribution of IRF8 to anti-tumor immunity, we implanted tumor cells into WT mice and mice harboring B cell-specific deletion of *Irf8*. Loss of IRF8 selectively in B cells was sufficient to significantly accelerate tumor growth (Fig. 6E), demonstrating that B cell-intrinsic IRF8 is required for effective tumor immune control.

### Pharmacologic activation of B cells restores anti-tumor immunity

Our above findings determined that loss of IRF8 was associated with impaired antigen presentation signatures, reduced plasmablast abundance, and defective immunostimulatory transcriptional programs in B lineage cells, suggesting that IRF8 is required for optimal B cell functional maturation. Because CD40 signaling is a central pathway regulating B cell activation and function ^47,48^, we hypothesized that pharmacologic activation of B cells using an agonistic anti-CD40 antibody could restore anti-tumor immune function in the setting of impaired IRF8-dependent B cell activation (Fig. S7A). To determine the translational relevance of this pathway in human tumors, we first analyzed publicly available human tumor scRNA-seq datasets. In human melanoma, CD40 expression was highly enriched in B cells, hypoxia-associated monocytes, monocyte/macrophage populations, and reactive microglia (Fig. S7B). Notably, clusters exhibiting high IRF8 expression also displayed elevated CD40 expression, suggesting coordinated regulation of immune activation programs (Fig. S7C). Similarly, scRNA-seq analyses of human colon tumors demonstrated strong CD40 expression in B cells, DCs, macrophages, monocytes, and plasma cells, with lower expression detected in subsets of T cells (Fig. S7D). Again, immune populations with high IRF8 expression showed correspondingly elevated CD40 expression (Fig. S7E), supporting the hypothesis that IRF8-associated immune activation states are linked to CD40 signaling pathways across human tumors.

We next tested whether pharmacologic CD40 activation could induce anti-tumor immune responses in vivo using an orthotopic CT26 colon tumor model (Fig. S7F). Tumor-bearing mice treated with agonistic anti-CD40 antibody developed marked splenomegaly relative to isotype-treated controls, consistent with systemic immune activation (Fig. S7G). Flow cytometric analyses revealed significant expansion of splenic CD8⁺ and CD4⁺ T cell populations following anti-CD40 treatment, whereas total CD19⁺ B cell numbers were not significantly altered (Fig. S7H-I). Importantly, anti-CD40 treatment also significantly increased tumor-infiltrating CD8⁺ T cells within colon tumors (Fig. S7J-K), indicating enhanced local anti-tumor immune activation. Consistent with these immunologic changes, anti-CD40 therapy significantly reduced tumor burden compared to control-treated mice (Fig. S7L). Taken together, these findings demonstrate that pharmacologic activation of B cells is sufficient to suppress tumor progression and further establish B cells as critical contributors to anti-tumor T cell immunity in the tumor microenvironment.

Taken together, these above results provide functional evidence that B cells contribute directly to T cell activation and anti-tumor immunity through their antigen-presenting function, and that this activity is governed intrinsically by IRF8. Combined with our transcriptomic and chromatin accessibility analyses, these findings support a model in which IRF8 establishes the immunostimulatory transcriptional and chromatin programs in B cells that are required for effective T cell activation and tumor immune control.

### IRF8-regulated B cell transcriptional programs are associated with response to PD-1 blockade immunotherapy in human cancer

To determine the human cancer relevance of the tumor-infiltrating B cell-intrinsic IRF8-regulated transcriptional program identified in our mouse tumor models, we first analyzed a published pre-treatment single-cell RNA-Seq dataset from breast cancer patients treated with neoadjuvant pembrolizumab (GSE160246) ^49^, in which patients were classified as T cell clonotype expanders (E) or non-expanders (NE) based on complementary scTCR-seq annotations. Re-clustering of B cells identified seven subclusters, including memory B cells, plasmablasts, naive B cells, transitional B cells, plasma cells, Ig-high B cells, and proliferating B cells, which were distributed across both expander and non-expander samples (Fig. 7A). Quantitative analysis revealed that plasmablast proportions were significantly enriched in expanders relative to non-expanders, both as a fraction of total B cells and as a fraction of all immune cells (Fig. 7B), supporting a specific association between plasmablast abundance and effective anti-tumor immune responses.

**Figure 7.**
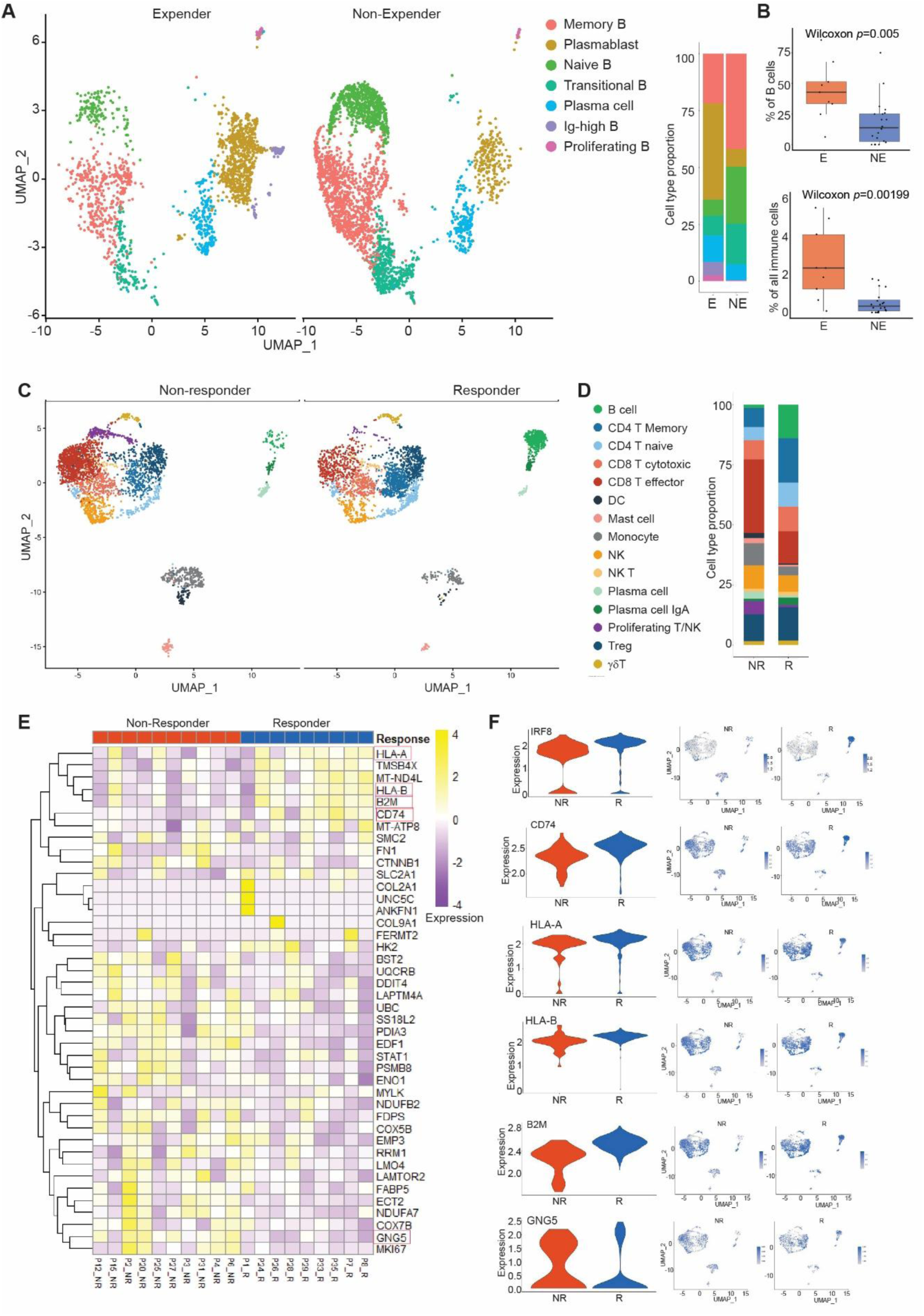
IRF8-regulated B cell transcriptional programs are associated with response to immunotherapy in human cancer. **A.** UMAP visualization of B cell subclusters identified in clonotype expanders (E) and non-expanders (NE) from pre-treatment scRNA-seq data of breast cancer patients treated with neoadjuvant pembrolizumab. scRNA-seq datasets were extracted from the GEO database (accession #GSE160246). **B.** Quantification of plasmablast proportions in expanders (E) versus non-expanders (NE), shown as percentage of total B cells. **C.** UMAP visualization of integrated scRNA-seq datasets showing immune cell populations identified in responders (R) and non-responders (NR) to PD-1 blockade immunotherapy in patients with melanoma. scRNA-Seq data were extracted from the GEO database (accession #GSE120575). **D.** Bar plot of cell type proportion as shown in C. **E.** Heatmap showing the expression of human orthologs of tumor-infiltrating mouse B cell-intrinsic IRF8-regulated genes in responders and non-responders to PD-1 blockade immunotherapy melanoma patients as shown in C. **F.** Violin plots (left) and UMAP feature plots (right) showing the expression levels of representative IRF8-regulated genes as indicated in non-responders (red) versus responders (blue) as shown in C.

We next analyzed a published single-cell RNA-seq dataset from melanoma patients treated with PD-1 blockade immunotherapy (GSE120575) ^50^. UMAP visualization of integrated responder and non-responder samples revealed preserved major immune cell populations but distinct differences in immune composition and activation states between clinical response groups (Fig. 7C). Quantitative analysis of immune cell composition further demonstrated enrichment of B cell-associated populations including plasma cells and plasmablasts in responders relative to non-responders (Fig. 7D), supporting a potential role for B cell immunostimulatory programs in mediating effective checkpoint blockade responses. We then converted the IRF8-regulated genes identified in mouse plasmablasts to their human orthologs. The IRF8-regulated B cell gene signature was significantly enriched in responders compared with non-responders in the melanoma cohort (Fig. 7E), suggesting that IRF8-dependent B cell programs are associated with favorable immunotherapy responses in human cancer. Consistent with the heatmap analysis, violin plot analyses demonstrated significantly changed expression of representative B cell-intrinsic IRF8-regulated genes, including IRF8, CD74, HLA-A, HLA-B, B2M, and GNG5, in responders relative to non-responders (Fig. 7F). These results support the human cancer relevance of our mouse findings and identify IRF8-associated B cell transcriptional programs as potential correlates of effective anti-tumor immunity and response to PD-1 blockade in human breast and melanoma.

## DISCUSSION

IRF8 has been established as a master regulator of myeloid cell differentiation and function, controlling dendritic cell development, restraining IMC accumulation, and regulating antigen cross-presentation and CD8⁺ T cell priming within tumors ^1,8,9,12,40,51,52^. IRF8 is also expressed throughout B cell development, where it governs germinal center formation, marginal zone versus follicular fate decisions, and plasma cell differentiation ^21,24,25,28,29,51,53^. We demonstrated in this study that IRF8 operates a B cell-intrinsic transcriptional program controlling antigen processing and presentation function to promote CD8⁺ T cell activation and tumor growth control. Critically, B cell-specific deletion of IRF8 was sufficient to significantly accelerate tumor growth, establishing a direct, cell-intrinsic requirement for IRF8 in B cell-mediated anti-tumor immunity independent of its broader myeloid functions. scRNA-seq differential expression analysis revealed that IRF8-deficient plasmablasts consistently downregulate MHC class I antigen presentation machinery genes, as well as interferon-response genes. Strikingly, the same antigen presentation gene module was coordinately suppressed in IRF8-deficient DCs and IMCs, with substantial overlap among the differentially expressed genes across all three populations. This cross-lineage conservation of an IRF8-dependent antigen presentation signature suggests that IRF8 orchestrates a conserved immunostimulatory module that extends well beyond canonical DC function to include B lineage cells. Integrated ATAC-RNA analyses demonstrated a strong positive correlation between IRF8-like peak accessibility loss and transcriptional downregulation of linked genes across all three populations. GO enrichment of genes linked to IRF8-like DA peaks identified antigen processing and presentation as the top enriched biological pathway in DCs, plasmablasts, and IMCs alike, directly connecting IRF8 enhancer accessibility to B cell antigen presentation function.

Prior work established that IRF8 establishes lineage-specific enhancer accessibility at AICE-containing distal elements in DCs and that IRF8-dependent chromatin opening precedes transcriptional activation in pDCs and microglia ^6,19,20,54^. We demonstrated here that plasmablasts in the TME exhibit an equally broad scope of IRF8-dependent chromatin accessibility regulation as in DCs and IMCs, with approximately 70% of differentially accessible peaks located in intronic and distal intergenic regions that constitute putative enhancers, a genomic distribution indistinguishable from that of DCs and IMCs. Motif enrichment analysis within differentially accessible regions identified 38 IRF8-like motifs spanning canonical ISRE, EICE, AICE, and PU.1/ETS composite elements across these myeloid and B cell lineages. However, lineage-specific motif preferences are distinct. ISRE motifs predominate in IMCs, a broad ETS/ISRE landscape characterizes DCs, whereas EICE composite elements are selectively enriched in plasmablasts, reflecting IRF8 cooperation with IRF4 and PU.1 in a B lineage-specific regulatory mode ^14^. These findings suggest that IRF8 engages distinct co-factor partners in each cell type to regulate overlapping but context-adapted antigen presentation programs. Notably, the functional consequences of IRF8 activity appear to be lineage- and context-dependent rather than uniformly immunostimulatory. In exhausted CD8^+^ T cells, IRF8 drive exhaustion-associated gene programs, such that IRF8 deficiency impairs exhausted T cell differentiation and restrains antitumor function ^38,55^. This contrasts with the antigen presentation-promoting, immunostimulatory role of IRF8 we observe in tumor-infiltrating B cells and the IRF8-DC axis in anti-cancer CD8^+^ T cell responses ^56^, suggesting that IRF8 does not have a fixed valence in anti-tumor immunity but instead executes distinct transcriptional programs depending on the chromatin and co-factor context of the cell in which it acts^52,57^.

The functional importance of B cell-intrinsic antigen presentation is directly demonstrated by our in vivo depletion, activation, and B cell-specific IRF8 deletion experiments. Antibody-mediated B cell depletion significantly accelerated tumor growth, hence establishing a non-redundant role for B cells in restraining tumor growth. Ex vivo co-culture experiments confirmed that B cells from tumor-bearing mice drive T cell proliferation in a ratio-dependent manner, directly demonstrating the T cell stimulatory capacity of B cells in tumor-bearing hosts. These findings are consistent with the established function of B cells as professional APCs that activate T cells through MHC-mediated antigen presentation and costimulatory molecule expression ^33,34,58,59^, and with the recent demonstration that blocking plasma cell differentiation enhances B cell antigen-presenting activity and anti-tumor immunity ^28^. Collectively, our findings establish that IRF8 preferentially engages EICE composite motifs at antigen presentation gene loci to act as a conserved epigenetic regulator of B cell antigen-presenting competence in the tumor microenvironment. Importantly, the heterogeneous roles ascribed to tumor-associated B cells in cancer, ranging from pro- to anti-tumorigenic, may reflect, in part, differences in B cell IRF8 activity and downstream antigen presentation competence ^29,60^. Lineage-specific loss or gain of IRF8-dependent transcriptional programs may thus serve as a molecular checkpoint controlling whether B cells adopt immunostimulatory or immunosuppressive functional states in the tumor microenvironment.

Analysis of published human cancer scRNA-seq datasets provides translational relevance for our mouse findings. In breast cancer scRNA-seq data from patients treated with neoadjuvant pembrolizumab, plasmablast proportions were significantly enriched in clonotype expanders relative to non-expanders, suggesting that IRF8-driven plasmablast identity correlates with productive anti-tumor immune responses. In melanoma patients treated with PD-1 blockade, human orthologs of the mouse IRF8-regulated B cell antigen presentation gene signature were significantly enriched in responders versus non-responders, including genes encoding MHC class I machinery components and interferon-response regulators. These results align with multiple clinical studies demonstrating that intratumoral B cells, plasma cells, and tertiary lymphoid structures associate with improved responses to immune checkpoint blockade across melanoma and other human cancers ^31,32,61,62^, and position IRF8 as a transcription factor that licenses these clinically relevant B cell immune states.

Beyond its established role in myeloid cells, our finding that IRF8 governs a conserved antigen-presentation program in tumor-infiltrating B cells suggests that therapeutic strategies currently designed to harness IRF8 in dendritic cells could be extended to the B lineage ^27,29,42^. LNP-encapsulated IRF8 mRNA has recently been shown to reprogram myeloid cells toward a cross-presenting, cDC1-like phenotype after both intratumoral and systemic delivery, generating durable antitumor immunity and synergizing with anti-PD-1 blockade in multiple syngeneic models ^42^. Given that B cells engage an analogous IRF8-dependent transcriptional module, favoring EICE-driven antigen-presentation gene expression over plasma cell differentiation, LNP-based delivery of IRF8 to tumor-infiltrating or tumor-draining B cells, potentially combined with strategies that restrain Blimp-1-driven plasma cell commitment, may represent a complementary route to expand the pool of antigen-presenting, costimulatory B cells competent to prime CD8⁺ T cells. This is particularly relevant for human translation since a constitutive interferon-high immunophenotype, marked by antigen-presentation gene induction (including CD74) in macrophages and tumor cells at sites of T cell infiltration has been shown to predict checkpoint inhibitor response in colorectal cancer independently of tumor mutational burden or mismatch-repair status^63,64^. Because B cells contribute to the same interferon-driven, antigen-presentation-competent tumor microenvironment, mRNA-LNP delivery of IRF8, engineered for lymphoid-organ tropism rather than hepatic uptake, could in principle convert CD74-low, immunotherapy-resistant tumors into CD74-high, ICI-responsive ones by simultaneously licensing antigen presentation in myeloid and B cell compartments ^63,65^, offering a tractable strategy to overcome resistance to immune checkpoint blockade^28^.

In summary, our findings extend IRF8 function beyond its canonical myeloid and DC roles, establish the tumor-infiltrating B cell as a critical IRF8-responsive antigen-presenting cell, and define B cell-intrinsic IRF8 expression as a transcriptional determinant of B cell immunostimulatory competence. The B cell-specific IRF8 deletion data directly demonstrate that this is a cell-autonomous function rather than a consequence of altered myeloid composition. Mechanistically, these results provide a framework for understanding how IRF8-dependent B cell differentiation state regulates the quality of anti-tumor T cell responses, and for identifying patients whose tumors harbor IRF8-competent B cell programs and who may therefore be more likely to benefit from B cell-activating immunotherapy strategies. Targeting IRF8-dependent pathways in B cells, including through CD40 agonism, restoring IRF8 expression ^66^ or approaches that preserve B cell antigen-presenting identity over plasma cell fate ^28^, may represent novel strategies to enhance anti-tumor immunity and improve outcomes to checkpoint blockade immunotherapy ^47,48,67^.

## METHODS

### Cell lines

CT26 and B16F10 were obtained from American Type Culture Collection (ATCC). Cell lines were treated with plasmocin (Invivogen, San Diego, CA) prior to animal studies and tested bi-monthly for mycoplasma. Cells were mycoplasma-free at time of experiments.

### Mice

BALB/c and C57BL6 mice were obtained from Charles River Laboratories at NCI Frederic (Frederick, MD). IRF8 knock out (IRF8 KO) mice were generated as previously described ^1^. Mice with the *loxp*-flanked *Irf8* gene [B6(Cg)-*Irf8^tm^*^1^*^.1Hm^*/J] were generated as previously described ^68^. Both male and female animals were used. Animals were age matched. Use of animal studies were approved by Augusta University (Protocol #2008–0162) and Charlie Norwood VA Medical Center Institutional Animal Care and Use Committees (Protocol #1314554–16).

### Tumors for Multiomics

B16-F10 cells (2.5×10^5^ cells/mouse in 100 μl PBS) were injected subcutaneously into WT or IRF8KO mice, one male and one female each for each mouse line. The tumors were dissected, processed with collagenase/hyaluronidase, filtered with 100 micron filter. Tumor infiltrating immune cells were isolated using a MojoSort CD45 selection kit (Cat# 480118, Biolegend) according to the manufacturer’s instructions.

### Tumor cell transplant

Tumor cells were cultured in RPMI+10%FBS for 24 hours, then harvested with trypsin and washed 3 times in PBS to remove media. Tumor cells were counted, viability assessed, and then resuspended in PBS for injection into R flank of mouse or into tip of cecum.

### In vivo B cell activation

Treatment began on days 3-10. Mice received 200ug of anti-CD40 or isotype control IgG in 100uL PBS intraperitoneal, every 3 days.

### Flow cytometry

Cells were resuspended as single cell suspension in cold PBS or FACS buffer. Antibodies were added at appropriate dilutions, and then stained for 30-60 minutes 4C, then washed with PBS+2% BSA. Cells were then fixed in 2% paraformaldehyde. Zombie UV Viability staining: Cells were resuspended as above. Zombie UV (1:1000) was added, and incubated for 10 minutes at room temperature, in the dark. Antibodies were then added at appropriate dilutions, incubated, washed, and fixed as above.

### Single-cell multiomics

Multiomics sequencing libraries including scRNA-seq and ATAC-seq libraries were generated using Chromium Next GEM Single Cell Multiome ATAC + 3’ Gene Expression Reagent Bundle, 4 rxns PN-1000285 (10x Genomics). Individual libraries were prepared for two independent biologic replicates containing WT and IRF8 KO mice with approximately 12,000 muclei on average per sample (total 24,000 cells each group). Cells were captured using the Chromium X (10x Genomics). The scRNAseq libraries were sequenced using Illumina Novaseq-6000 two SP runs, with a high output kit under the following sequencing protocol: 28 bp (Read 1), 10 bp, 10 bp (dual indexing Runs), and 90 bp (RNA Read 2) having Q30 bases in RNA read greater than 86% to collect approximately 16 K mean pre-normalization high-quality read pairs per cell, at the range of 1,700-3,900 median genes per cell, the ATAC-seq were sequenced using Illumina Novaseq-6000 one SP and one S2 runs with a high output kit under the following sequencing protocol: 50 bp (Read 1), 8 bp, 24 bp (dual indexing Runs), and 49 bp (RNA Read 2) having Q30 bases greater than 91% to collect approximately 72 K mean pre-normalization high-quality read pairs per cell, with the enrichment scores above 8.1. The raw reads in fastq format were processed using 10x Genomics Cell Ranger (7.0.0) analysis pipeline via STAR v2.5.0 alignment against murine mm10 reference genome. The Cell Ranger outputs containing gene-by-cell differential expression data, read counts, and normalized mapped reads aggregated among samples were imported into Loupe Browser (10x Genomics) for the cell marker annotations to further analyze cell populations in clusters of UMAP and TSNE plots.

### Single-Cell ATAC-seq data Processing and Chromatin Accessibility Analysis

Single-cell ATAC-seq data were processed using ArchR (v1.0.2) with the mm10 (GRCm38) reference genome. Arrow files were generated from fragment files for four samples [two IRF8 KO and two wild-type (WT) replicates]. An ArchRProject was constructed and cell barcodes were matched to a pre-processed Seurat multiome object containing dendritic cells (DCs), plasmablasts, and IMCs by extracting core barcode sequences (stripping sample prefixes and suffixes). Cell type and genotype metadata were transferred from the Seurat object to the ArchR project. Dimensionality reduction was performed using iterative latent semantic indexing (IterativeLSI) on the 500-bp tile matrix (25,000 variable features, 30 LSI dimensions, 2 iterations). Cells were clustered using the Seurat graph-based algorithm (resolution = 0.8) and visualized using UMAP (30 neighbors, minimum distance = 0.4). Pseudo-bulk ATAC-seq coverage tracks were computed per cell-type-condition group (ct_cond), and a reproducible peak set was called using MACS2 (v2.2.7) with default parameters. A peak-by-cell accessibility matrix (PeakMatrix) was subsequently added to the ArchR project.

### Differential Chromatin Accessibility Analysis

Differential accessibility (DA) analysis was performed independently for each cell type (DC, plasmablast, IMCs using per-cell-type ArchR sub-projects. For each cell type, chromatin accessibility was compared between irf8KO and WT cells using a binomial test on binarized peak accessibility values (getMarkerFeatures, testMethod = "binomial", binarize = TRUE). DA peaks were defined by a false discovery rate (FDR) < 0.01 and log₂ fold change (log₂FC) ≥1. Peaks with log₂FC>1 were designated as "gained" (increased accessibility in irf8KO), and peaks with log₂FC <-1 were designated as "lost" (decreased accessibility in irf8KO). Results were visualized as MA plots and volcano plots, in which the x-axis represents log₁₀(mean accessibility + 1) or log₂FC, respectively, and the y-axis represents log₂FC or -log₁₀(FDR).

### Identification of IRF8-like Motif-Containing Peaks

A curated repository of 38 IRF8-like position frequency matrices was loaded as a PFMatrixList object (TFBSTools). PFMs were converted to position weight matrices (PWMs) using toPWM() and added to the ArchR project as a custom motif annotation (addMotifAnnotations, cutOff = 1e-4). In parallel, motif scanning across all called peaks was performed using matchMotifs() from the motifmatchr package against the mm10 genome sequence (BSgenome.Mmusculus.UCSC.mm10). A peak was designated as "IRF8-like" if it contained at least one match to any of the 38 IRF8-like PWMs. Per-peak IRF8-like flags were saved as a reference table and subsequently merged with DA results by coordinate-based peak identifiers (chr:start-end).

### Genomic Annotation and Distribution Analysis

Genomic annotation of peak sets was performed using ChIPseeker (v1.36) with the mm10 gene annotation database (TxDb.Mmusculus.UCSC.mm10.knownGene). Promoter regions were defined as ±3 kb from the transcription start site (TSS). Annotation categories (e.g., promoter, intron, exon, intergenic, downstream) were simplified by removing sub-category parenthetical descriptors. For each cell-type-condition group, the fraction and count of IRF8-like peaks falling within each genomic category were tabulated and visualized as stacked bar charts and faceted bar plots. The genomic distribution of IRF8-like DA peaks (gained and lost in irf8KO) was similarly annotated and summarized across cell types.

### IRF8-like Motif Enrichment Analysis in Gained and Lost DA Peaks

For each cell type and DA direction (gained or lost in irf8KO), enrichment of individual IRF8-like motifs within the DA peak subset relative to a background comprising all accessible IRF8-like peaks from that cell type was assessed by two-sided Fisher’s exact tests. For each motif m and peak subset T, a 2×2 contingency table was constructed from the counts of motif-positive and motif-negative peaks in T versus the background. P-values were corrected for multiple testing using the Benjamini-Hochberg procedure. Motifs were considered significantly enriched at FDR < 0.05 and odds ratio > 1. Enrichment results were visualized as a combined heatmap (ComplexHeatmap) in which rows represent the 38 IRF8-like motif variants, columns represent cell-type-DA-direction groups, and color encodes −log₁₀(FDR), masked to zero for motifs with odds ratio ≤ 1. Motif sequence logos were rendered in information-content-scaled bit representation using the universalmotif package (view_motifs, use.type=ICM).

### Inter-peak Distance Analysis

To assess spatial clustering of IRF8-like peaks across the genome, inter-peak distances were computed for each cell-type-condition group. IRF8-like peaks assigned to each group (based on the MACS2 peak group label) were sorted by chromosomal coordinate, and the gap distance between the end of each peak and the start of the immediately downstream peak on the same chromosome was calculated. Distance distributions were compared between WT and irf8KO conditions for each cell type using kernel density estimation and visualized on a log₁₀ scale.

### Integration of Differential Chromatin Accessibility and Differential Gene Expression

Differential gene expression (DE) analysis between IRF8 KO and WT cells was performed for each cell type using Seurat’s FindMarkers function (Wilcoxon rank-sum test, minimum cell fraction = 0.1) on log-normalized RNA counts. DE genes were defined as FDR≤0.01 and log₂FC≥0.58 (∼1.5-fold). Peak-to-gene linkages were computed within each cell-type ArchR sub-project using addPeak2GeneLinks(IterativeLSI reduced dimensions, GeneScoreMatrix, maximum distance =100 kb) and filtered at Pearson correlation ≥ 0.45 and FDR ≤ 0.05. DA peaks were linked to genes using these peak-to-gene links, and the resulting gene sets were intersected with DE genes to identify concordantly altered loci. IRF8-like DA peaks were further restricted to those whose nearest annotated gene (TxDb.Mmusculus.UCSC.mm10.knownGene, via nearest()) was among the DE genes. Heatmaps displaying ATAC-seq accessibility profiles centered on IRF8-like DA peak summits (±1 kb, 10-bp bins) were generated using EnrichedHeatmap, with peaks split into "Up" and "Down" groups and ranked within each group by the magnitude of paired RNA expression difference (KO − WT average log-normalized counts) of the nearest DE gene.

### Gene Ontology Enrichment Analysis

For genes linked to IRF8-like DA peaks through either nearest-gene assignment or peak-to-gene linkage, Gene Ontology (GO) Biological Process enrichment analysis was performed using clusterProfiler (v4.8) with the enrichGO function (organism annotation: org.Mm.eg.db; gene ID type: Entrez; correction method: Benjamini-Hochberg; P-value cutoff: 0.05; Q-value cutoff: 0.20). Gene symbols were converted to Entrez identifiers using bitr(). The top 10 enriched GO terms per cell type were selected by adjusted P-value and visualized as dot plots and bar plots.

### Genome Browser Track Visualization

Genome browser-style coverage tracks for selected IRF8-like DA peaks and their nearest DE genes were generated using Signac’s CoveragePlot function applied to the multiome Seurat/Signac object, displaying pseudo-bulk ATAC-seq accessibility stratified by genotype (WT vs. irf8KO) over a ±25-kb window centered on each DA peak summit. Paired RNA expression levels of the nearest DE gene were overlaid as violin plots using VlnPlot. Regions lacking called peaks in the ATAC assay were excluded from visualization.

### Software and Statistical Analysis

All analyses were conducted in R (v4.3). Key packages used include ArchR (v1.0.2), Seurat (v5), Signac, motifmatchr, TFBSTools, universalmotif, ChIPseeker, clusterProfiler, EnrichedHeatmap, and ComplexHeatmap. Multiple testing correction was applied using the Benjamini-Hochberg method throughout. Figures were generated using ggplot2, with classic theme formatting applied globally.

### Generation of an IRF8-like motif atlas and integration of chromatin accessibility with single-cell transcriptomics

To define a reference set of IRF8-like motifs for downstream IRF8 peak annotation and motif enrichment analyses, we generated a curated IRF8 motif repository. Position frequency matrices (PFMs) were retrieved from the JASPAR 2022 database using TFBSTools, and additional IRF-related motifs from HOMER were incorporated when available. From the combined motif repository, we extracted an IRF-like subset based on motif IDs and names using a permissive literature-guided pattern that included canonical IRF8 and related IRF/ETS composite motifs, including *IRF8, IRF, ISRE, EICE, IECS, AICE, SPI1/PU.1,* and *ETS* family elements (references). This strategy was designed to capture canonical IRF8-binding motifs (e.g., MA0652.1), ISRE-like motifs, and composite IRF-ETS regulatory elements known to mediate IRF8-dependent transcriptional programs. The resulting curated IRF8-like motif set was subsequently used for all downstream analyses, including IRF8-like peak definition, motif scanning, motif enrichment, and sequence logo generation. Using this approach, we identified a total of 38 enriched IRF8-like motifs across the analyzed immune populations. To integrate chromatin accessibility with transcriptomic profiles, we matched cells between ArchR and Seurat by extracting shared core cellular barcodes and intersecting overlapping cell identities between datasets. The ArchRProject was then subset to matched cells and annotated using integrated metadata labels corresponding to cell type and genotype. This unified multiomic object served as the foundation for all subsequent chromatin accessibility and peak-to-gene linkage analyses.

### ArchR cell clustering validates immune cell identity and genotype separation

Iterative Latent Semantic Indexing (IterativeLSI) was performed on the TileMatrix of the ArchR project subset using 25,000 variable features and dimensions 1-30 to cluster cells in LSI space at a resolution of 0.8. Cells were subsequently embedded into two-dimensional space using UMAP based on LSI coordinates. The resulting clustering analysis resolved the major immune populations, including DCs, IMCs, and plasmablasts, confirming concordance between chromatin accessibility-based clustering and transcriptomic cell annotations. In addition, WT and Irf8-deficient cells exhibited distinct yet partially overlapping chromatin accessibility landscapes within each immune lineage, consistent with genotype-dependent epigenetic remodeling while preserving overall cell identity (Fig. 4A-B).

### Group coverage generation, peak calling, peak matrix construction, peak-to-gene linkage, and differential accessibility analysis

To characterize chromatin accessibility landscapes across immune populations and genotypes, scATAC-seq analyses were performed using the ArchR framework on the integrated multiomic dataset. Cells were grouped according to combined cell type and genotype annotations (ct_cond) to enable parallel analysis across immune lineages and experimental conditions. Group-level pseudo-bulk chromatin accessibility tracks were first generated using the addGroupCoverages() function in ArchR. These pseudo-bulk profiles were subsequently used for reproducible peak calling using addReproduciblePeakSet() with MACS2, generating a unified and reproducible consensus peak set across all cell type × genotype groups. This approach minimizes sparsity inherent to single-cell chromatin accessibility datasets while enabling robust identification of accessible regulatory regions shared across biologically relevant populations. Following peak calling, a peak-by-cell accessibility matrix (PeakMatrix) was constructed using addPeakMatrix(), producing a quantitative accessibility matrix for all cells across the unified peak set. The resulting PeakMatrix served as the foundation for downstream differential accessibility, motif enrichment, and peak-to-gene linkage analyses.

To associate distal regulatory elements with putative target genes, peak-to-gene linkage analysis was performed on the integrated ArchRProject_subset using addPeak2GeneLinks() with the GeneScoreMatrix and a maximum genomic linkage distance of 100 kb. This analysis identifies correlated chromatin accessibility and gene activity relationships between accessible peaks and nearby genes. Peak-to-gene associations were subsequently filtered and intersected with IRF8-like motif-containing peaks to identify candidate IRF8-regulated enhancer/promoter-gene interactions. For cell type-specific differential accessibility (DA) analyses, the integrated ArchR project was subset independently into DC, plasmablast, and IMC populations. Within each cell type-specific project, DA analysis between WT and Irf8 KO cells was performed on the PeakMatrix using a binomial test on binarized peak accessibility values implemented in ArchR. Bidirectional comparisons were conducted, including KO versus WT (peaks gained in KO) and WT versus KO (peaks lost in KO). Differentially accessible peaks were defined using thresholds of false discovery rate (FDR) ≤ 0.01 and absolute log2 fold-change≥1.

DA peak tables were exported for downstream analyses and visualization. MA plots and volcano plots were generated to visualize global genotype-dependent chromatin accessibility changes within each immune population. Genomic annotation analyses were performed to determine the distribution of DA peaks across promoter, intronic, exonic, and distal intergenic regions. In addition, bar plots comparing the number of peaks gained versus lost in Irf8-deficient cells were generated for each cell type. To infer biological pathways associated with IRF8-dependent chromatin remodeling, Gene Ontology (GO) enrichment analyses were performed on genes linked to DA peaks using peak-to-gene associations.

### Definition and genome-wide characterization of IRF8-like peaks

To identify and annotate IRF8-like regulatory elements genome-wide, we utilized a curated collection of 38 IRF8-like position frequency matrices (PFMs) representing canonical IRF8 motifs and related IRF/ETS composite motifs. PFMs were converted to position weight matrices (PWMs) and incorporated as a custom motif annotation set within the ArchR peak set using motif matching with a significance threshold of 1 × 10⁻⁴. Motif scanning generated, for each accessible peak, a binary motif match matrix indicating the presence or absence of each of the 38 IRF8-like motifs. An “IRF8-like peak” was defined as any chromatin accessibility peak containing at least one IRF8-like motif match. To focus on biologically relevant regulatory elements, analyses were restricted to IRF8-like peaks that were accessible within the corresponding cell type and genotype group based on the PeakMatrix. This filtering strategy enabled identification of active IRF8-associated regulatory regions utilized in each immune population under WT and Irf8-deficient conditions. Genome-wide annotation of accessible IRF8-like peaks was performed using the annotatePeak function with the TxDb.Mmusculus.UCSC.mm10.knownGene transcript database and a transcription start site (TSS) annotation window of ±3 kb. Peaks were classified into genomic categories including promoter, 5′ untranslated region (UTR), exon, intron, downstream, and distal intergenic regions. For each cell type and genotype, the fraction and total number of IRF8-like peaks within each genomic category were quantified. Data visualization included stacked bar plots showing proportional genomic distributions, faceted bar plots displaying peak counts by genomic annotation, and side-by-side WT versus Irf8 KO comparison plots to assess genotype-dependent shifts in IRF8-associated chromatin accessibility landscapes.

### Human cancer patient scRNA-seq dataset analysis

To assess the clinical relevance of IRF8-dependent plasmablast differentiation, we analyzed pre-treatment scRNA-seq data from 29 breast cancer patients treated with neoadjuvant pembrolizumab (Cohort 1: GSE160246). Patients were classified as clonotype expanders (E, n=9) or non-expanders (NE, n=20) based on complementary scTCR-seq annotations. B cells were re-clustered in Seurat (resolution 0.3) and subclusters were annotated by marker gene expression. Plasmablasts were identified by the expression of PRDM1, IGHG4, and immunoglobulin light chain genes; contaminating T cell and epithelial clusters were excluded. Plasmablast proportions relative to total immune cells were compared between E and NE patients by Wilcoxon rank-sum test.

### Data analysis

Statistical analysis was done using Graphpad Prism V10.0 and one way ANOVA with Tukey’s multiple comparisons and paired Student’s t-test to determine statistical significance. A p<0.05 is considered as significant in difference.

## Acknowledgements

We acknowledge the support and contribution of the Integrated Genomics Core Shared Resources at the Georgia Cancer Center at Augusta University (RRID: SCR_026483).

## Author Contributions

Conceptualization (ZT, DBP, HS, KS, KO, and KL), Methodology: (ZT, DBP, RR, SB, KF, PC, DY), Funding acquisition: (KL), Project administration (KL), Bioinformatics (RR, SB, HS), Writing: original draft (ZT, DBP, KL), Writing: review & editing: (ZT, DBP, KL)

## Conflict of interest

None.

## Funding

Grant support from the US Department of Veterans Affairs I01CX001364 (to K.L). National Cancer Institute R01CA278852 (to K.L.), and R43CA287611 (to P.S.R.).

## Data availability

The single-cell multiomics datasets are deposited in GEO with accession #GSE334958.

## Notes

Conflict of interest: The authors declare no potential conflicts of interest

### Competing Interest Statement

The authors have declared no competing interest.

