## Supplemental Figures and tables for "B cell-intrinsic IRF8 transcriptionally reprograms antigen presentation to sustain CD8⁺ T cell antitumor immunity"

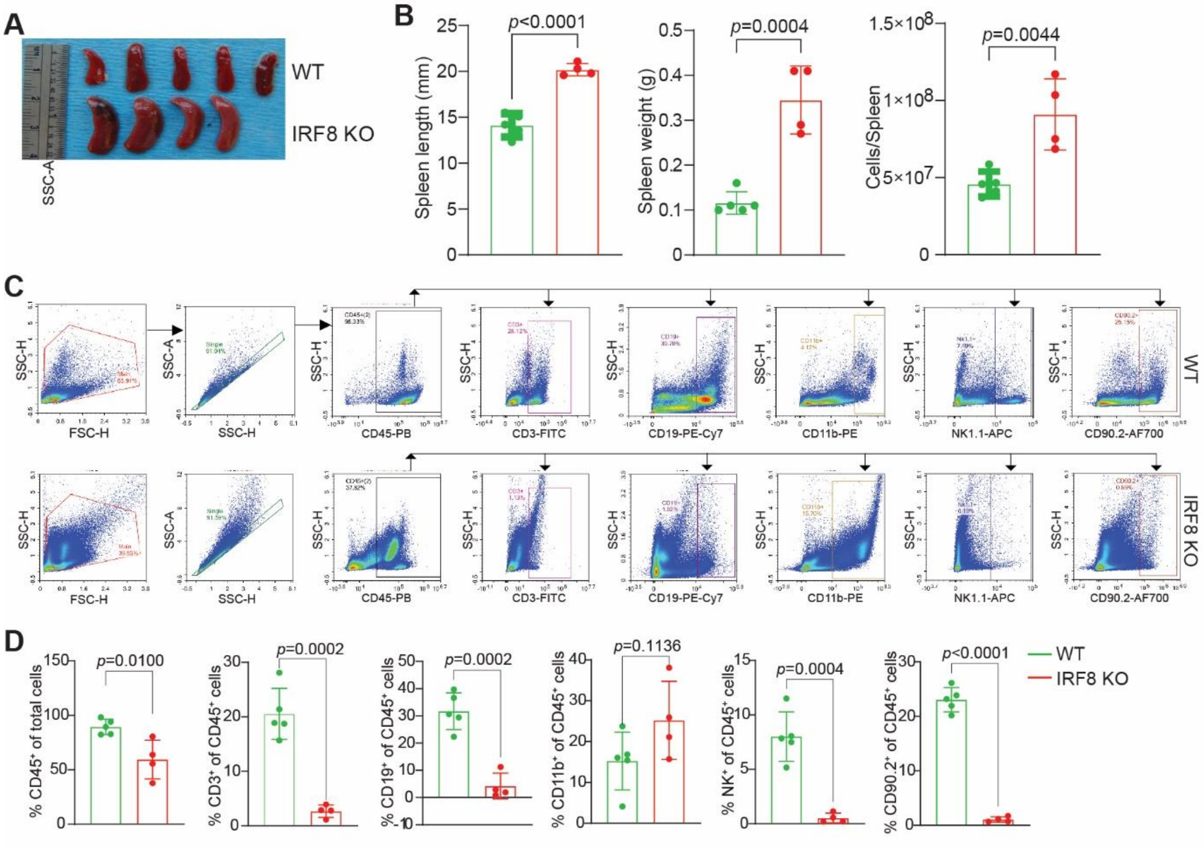

**Figure S1. IRF8 deficiency causes splenomegaly and reduced lymphoid representation in tumor-bearing mice.**

**A.** Images of spleens isolated from B16-F10 tumor-bearing WT and IRF8 KO mice.

**B.** Quantification of spleen length, weight, and total splenocyte number in WT and IRF8 KO tumor-bearing mice.

**C.** Flow cytometry gating strategy for splenic cells in WT (top) and IRF8 KO (bottom) tumor-bearing mice.

**D.** Quantification of the indicated cell types in tumor-bearing mouse spleens. Data are presented as mean ± SD.

DotPlot GSEA GOBP PlasmaBlast

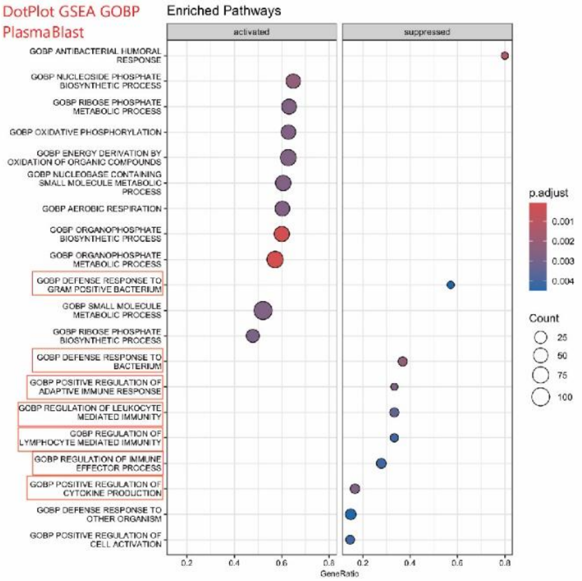

DotPlot GSEA GOBP DC

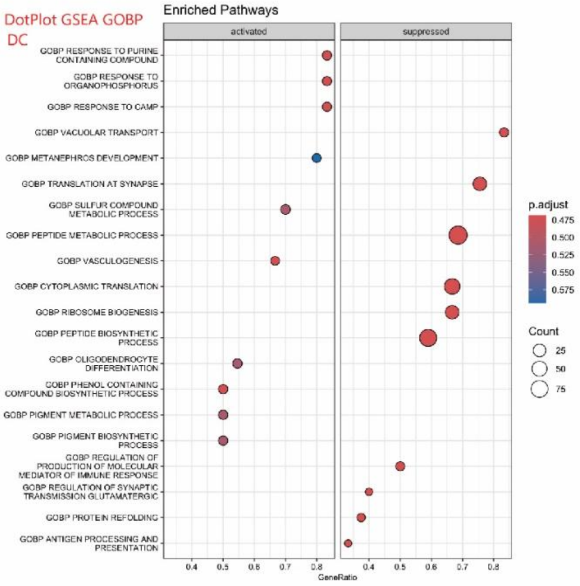

DotPlot GSEA GOBP IMC

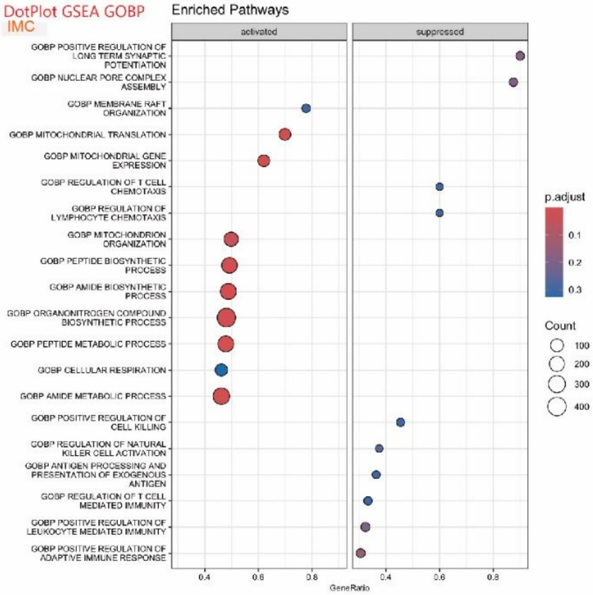

**Figure S2. Gene Ontology enrichment analyses of IRF8-dependent transcriptional programs in tumor-infiltrating immune populations.** Dot plot Gene Ontology Biological Process (GOBP) enrichment analyses of differentially expressed genes in plasmablasts (top), dendritic cells (DCs; middle), and MDSCs (bottom) isolated from WT and Irf8 KO tumors. Enriched pathways are separated into activated and suppressed transcriptional programs based on differential expression patterns between WT and IRF8 KO cells. Dot size represents the number of genes associated with each pathway, whereas color intensity indicates adjusted P values.

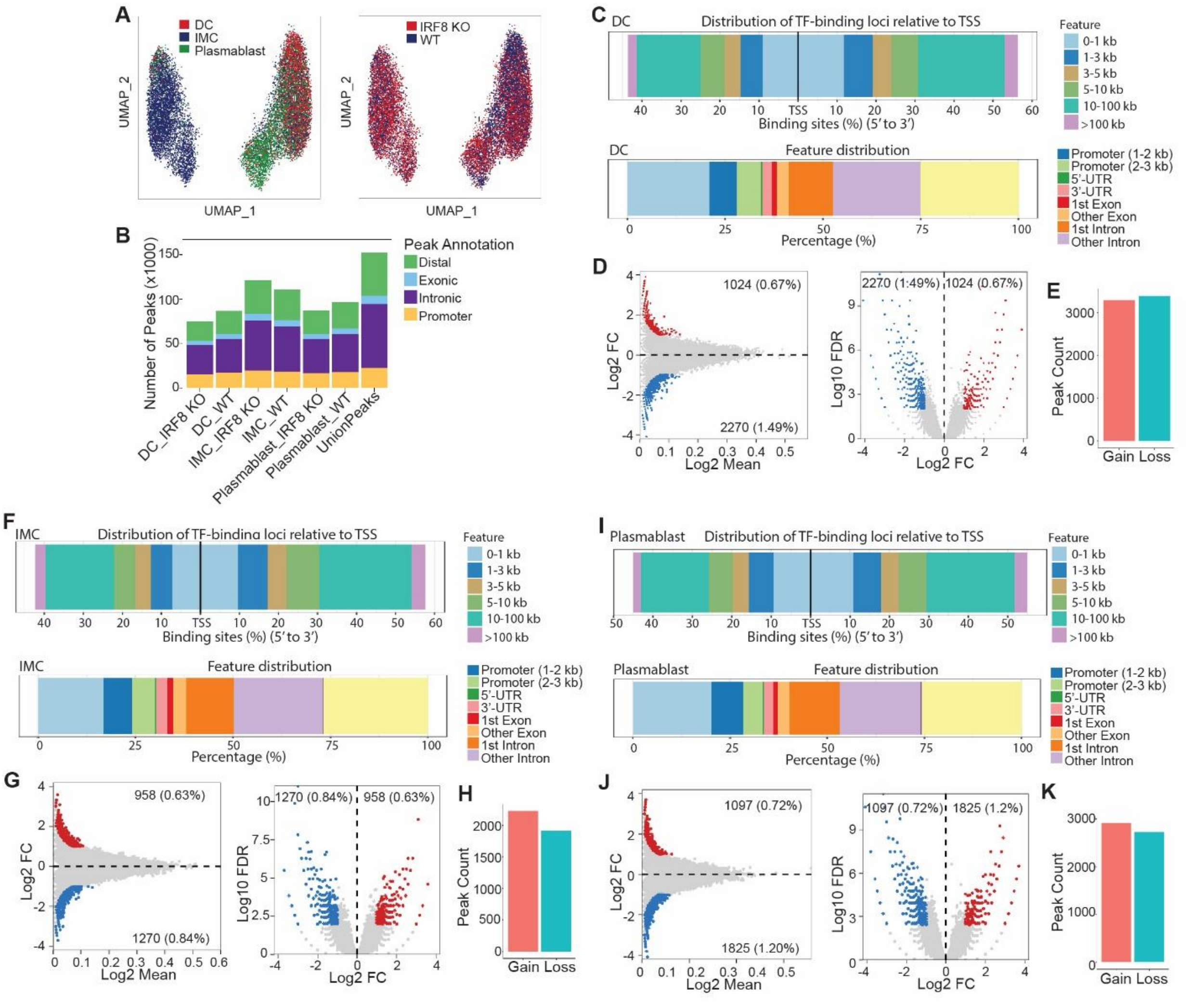

**Figure S3. Single-cell ATAC-seq analysis reveals IRF8-dependent chromatin accessibility remodeling across myeloid and B lineage cells.**

A. UMAP visualization of integrated scATAC-seq profiles of tumor-infiltrating immune cell types as indicated from B16-F10 tumor-bearing WT and IRF8 KO mice following Iterative LSI dimensionality reduction (25,000 variable features, dimensions 1-30) and

UMAP embedding (nNeighbors=30, minDist=0.4), colored by immune cell types (left) and genotype (right).

B. Genomic annotation of accessible chromatin peaks containing at least one IRF8-like motif match (n=38-motif curated PFM set; motifmatchr, mm10 genome) across immune cell types and genotypes, showing the proportional distribution within promoter (≤1 kb, 1-2 kb, and 2-3 kb), exon, intron, and distal intergenic regions (ChIPseeker; TxDb.Mmusculus.UCSC.mm10.knownGene; TSS±3kb).
C. Distribution of differentially accessible (DA) chromatin peaks relative to TSSs in DCs, plotted as the proportion of peaks at varying distances from TSSs (binomial DA test, binarized accessibility; FDR≤ 0.01, Log2FC≥ 1). Top panel: binding site distribution; bottom panel: genomic feature distribution.
D. Mean-average (MA; left) and volcano (right) plots of DA peaks comparing WT and IRF8 KO DCs across all accessible peaks. Grey points: non-significant peaks; red: IRF8-like motif-containing peaks with significantly increased accessibility in IRF8 KO (Log2FC≥ 1, FDR≤0.01); blue: IRF8-like motif-containing peaks with significantly decreased accessibility in Irf8 KO (Log2FC≤−1, FDR≤0.01). Peak counts and percentages of total are annotated per quadrant. Dashed lines indicate DA thresholds (Log2FC=1; FDR=0.01).
E. Quantification of chromatin accessibility peaks gained (increased in IRF8 KO) or lost (decreased in IRF8 KO) in DCs relative to WT controls (FD ≤0.01, Log2FC≥1; binomial test on binarized PeakMatrix).
F. Distribution of DA chromatin peaks relative to TSSs in MDSCs (same parameters as C).
G. MA (left) and volcano (right) plots of DA peaks comparing WT and IRF8 KO MDSCs. Color scheme and thresholds as in (D).
H. Quantification of chromatin accessibility peaks gained or lost in IRF8 KO MDSCs relative to WT controls.
I. Distribution of DA chromatin peaks relative to TSSs in plasmablasts (same parameters as C).
J. MA (left) and volcano (right) plots of DA peaks comparing WT and Irf8 KO plasmablasts. Color scheme and thresholds as in (D). Numbers in quadrants indicate count and percentage of significantly altered IRF8-like motif-containing peaks relative to all evaluated peaks.
K. Quantification of chromatin accessibility peaks gained or lost in IRF8 KO plasmablasts relative to WT controls. DA analysis was performed using the binomial test on binarized per-cell accessibility scores (ArchR; FDR≤0.01, Log2FC≥1).

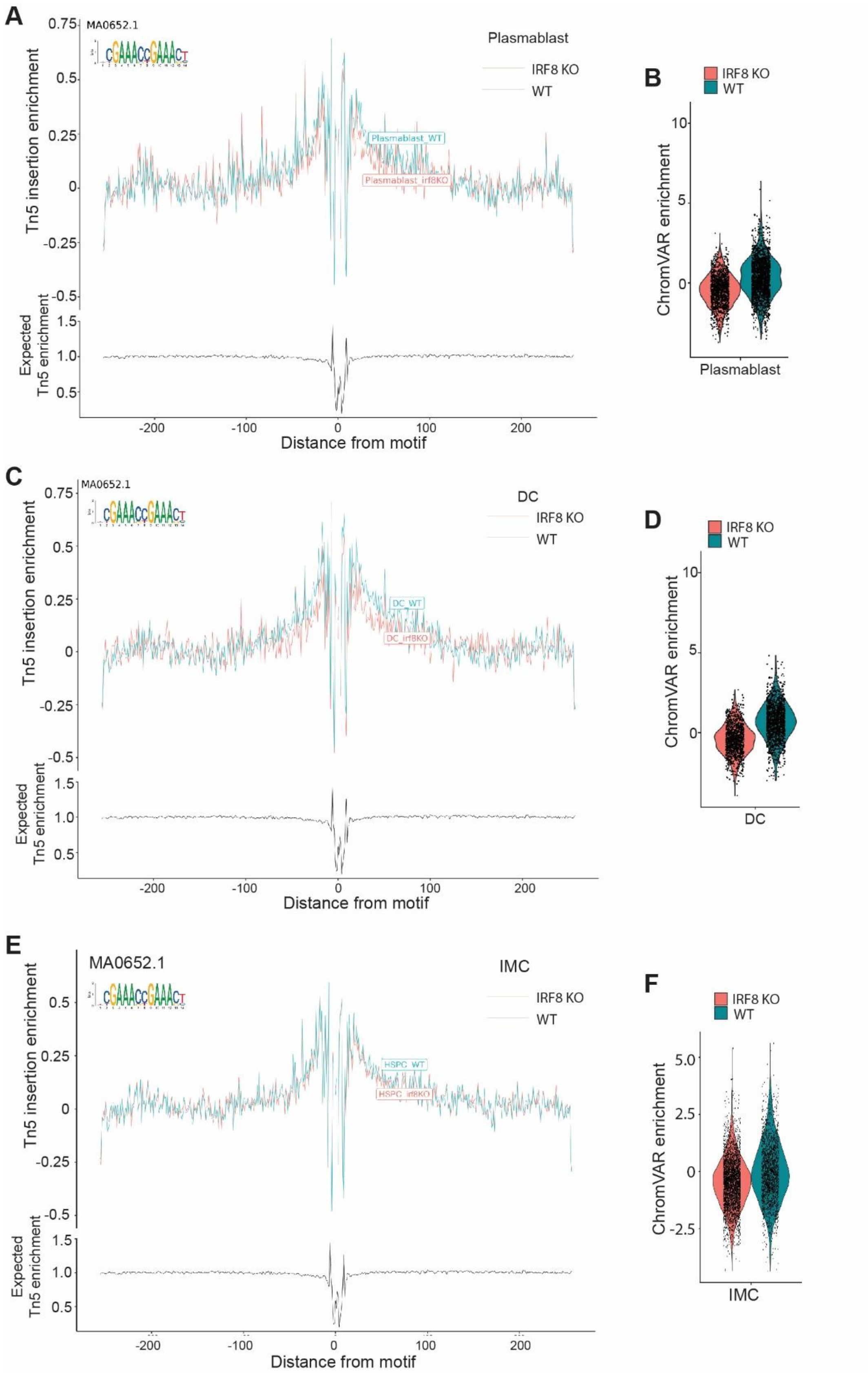
A
MA0652.1
Plasmablast
IRF8 KO
WT
Plasmablast_WT
Plasmablast_irf8KO
Tn5 insertion enrichment
0.75
0.5
0.25
0
-0.25
-0.5
Expected
Tn5 enrichment
1.5
1.0
0.5
-200
-100
0
100
200
Distance from motif
B
IRF8 KO
WT
ChromVAR enrichment
10
5
0
Plasmablast
C
MA0652.1
DC
IRF8 KO
WT
DC_WT
DC_irf8KO
Tn5 insertion enrichment
Expected
Tn5 enrichment
Distance from motif
D
IRF8 KO
WT
ChromVAR enrichment
DC
E
MA0652.1
IMC
IRF8 KO
WT
HSPC_WT
HSPC_irf8KO
Tn5 insertion enrichment
Expected
Tn5 enrichment
Distance from motif
F
IRF8 KO
WT
ChromVAR enrichment
5.0
2.5
0
-2.5
IMC

**Figure S4. IRF8 motif footprinting and accessibility are reduced at IRF8-like loci in IRF8 KO plasmablasts and DCs.**

**A**. Tn5 insertion enrichment (footprinting) profile centered on the canonical IRF8 motif (MA0652.1) in WT (blue) and IRF8 KO (orange) plasmablasts. Top: Tn5 insertion enrichment as a function of distance from the motif center. Bottom: corresponding expected Tn5 enrichment.

**B**. Violin plot of IRF8 motif accessibility scores in WT and IRF8 KO plasmablasts as shown in A.

**C**. Tn5 insertion enrichment (footprinting) profile centered on the canonical IRF8 motif (MA0652.1) in WT (blue) and IRF8 KO (pink) DCs, shown as in panel A.

D. Violin plot of IRF8 motif accessibility scores in WT and IRF8 KO DCs as in C.

**E**. Tn5 insertion enrichment (footprinting) profile centered on the canonical IRF8 motif (MA0652.1) in WT (blue) and IRF8 KO (pink) MDSCs, shown as in panel A.

F. Violin plot of IRF8 motif accessibility scores in WT and IRF8 KO MDSCss as in E.

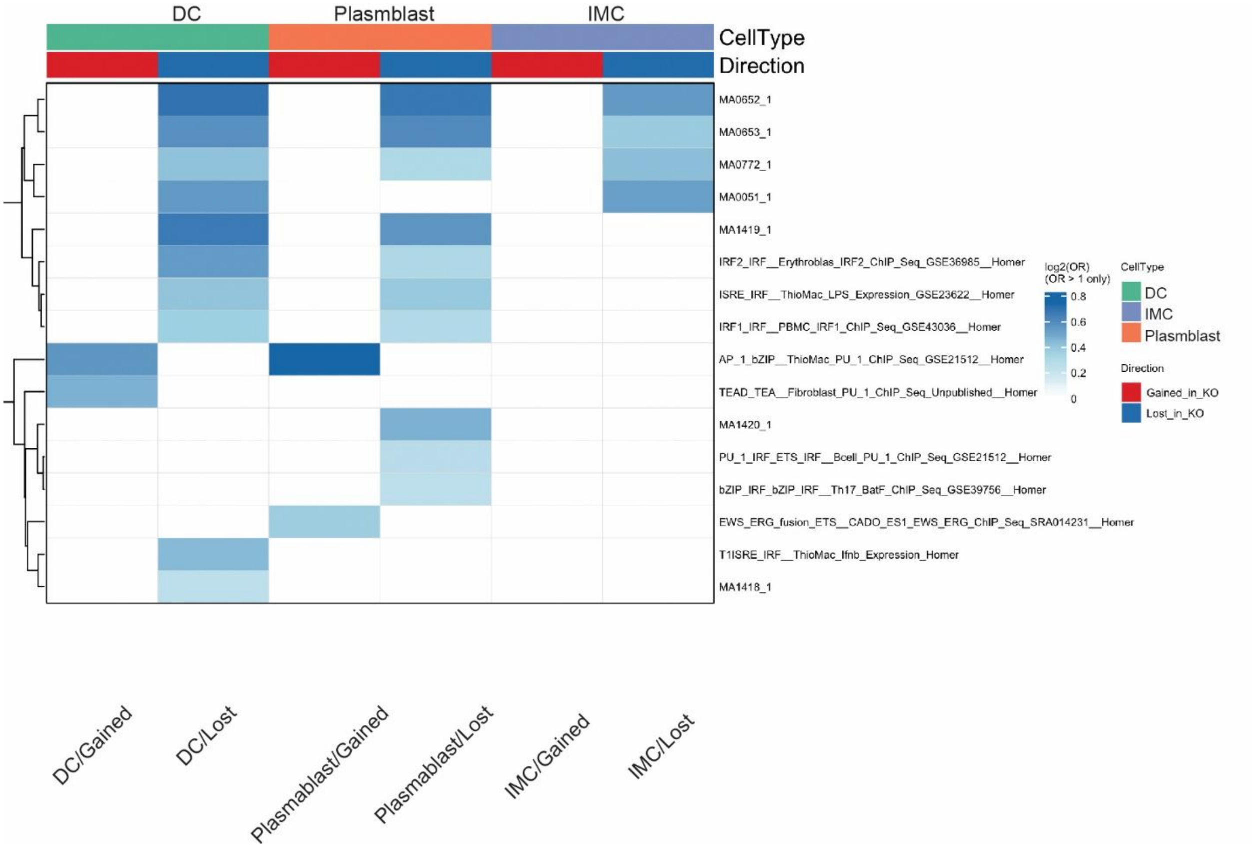

**Figure S5. IRF8-dependent transcription factor motif remodeling across tumor-infiltrating immune populations**. Shown is heatmap of enrichment of IRF8-like motifs identified from integrated single-cell ATAC-seq and scRNA-seq analyses of DCs, plasmablasts, and MDSCs from WT and IRF8 KO mice. DA chromatin regions identified by scATAC-seq were integrated with corresponding transcriptional programs from matched scRNA-seq datasets to define IRF8-associated regulatory architectures across immune lineages. Columns represent immune cell populations and chromatin accessibility states (peaks gained or lost in IRF8 KO mice relative to WT), whereas rows represent enriched IRF8-associated motifs.

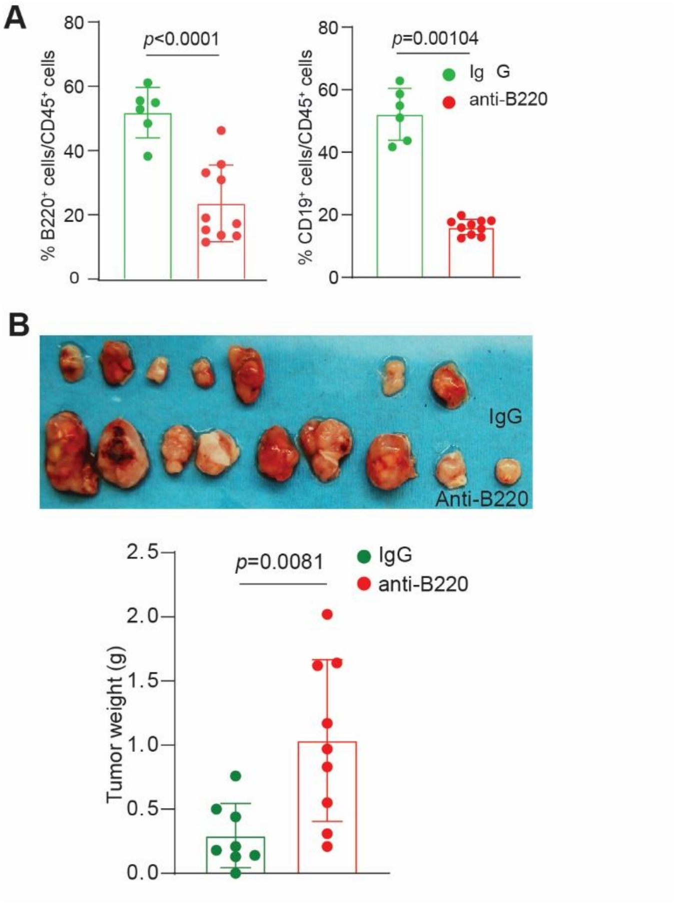

**Figure S6. B220 depletion accelerates tumor growth in mice.**

**A.** CT26 tumor-bearing mice were treated with B220 neutralization mAb. Shown is quantification of the percentage of $B220^+$ and $CD19^+$ cells among $CD45^+$ splenocytes from mice treated with control IgG or B220-neutralizing monoclonal antibody.

**B.** Top: Images of CT26 tumors from IgG- and anti-B220-treated mice. Bottom: Quantification of tumor weights at experimental endpoint in IgG and anti-B220-treated mice. Data are presented as mean ± SD.

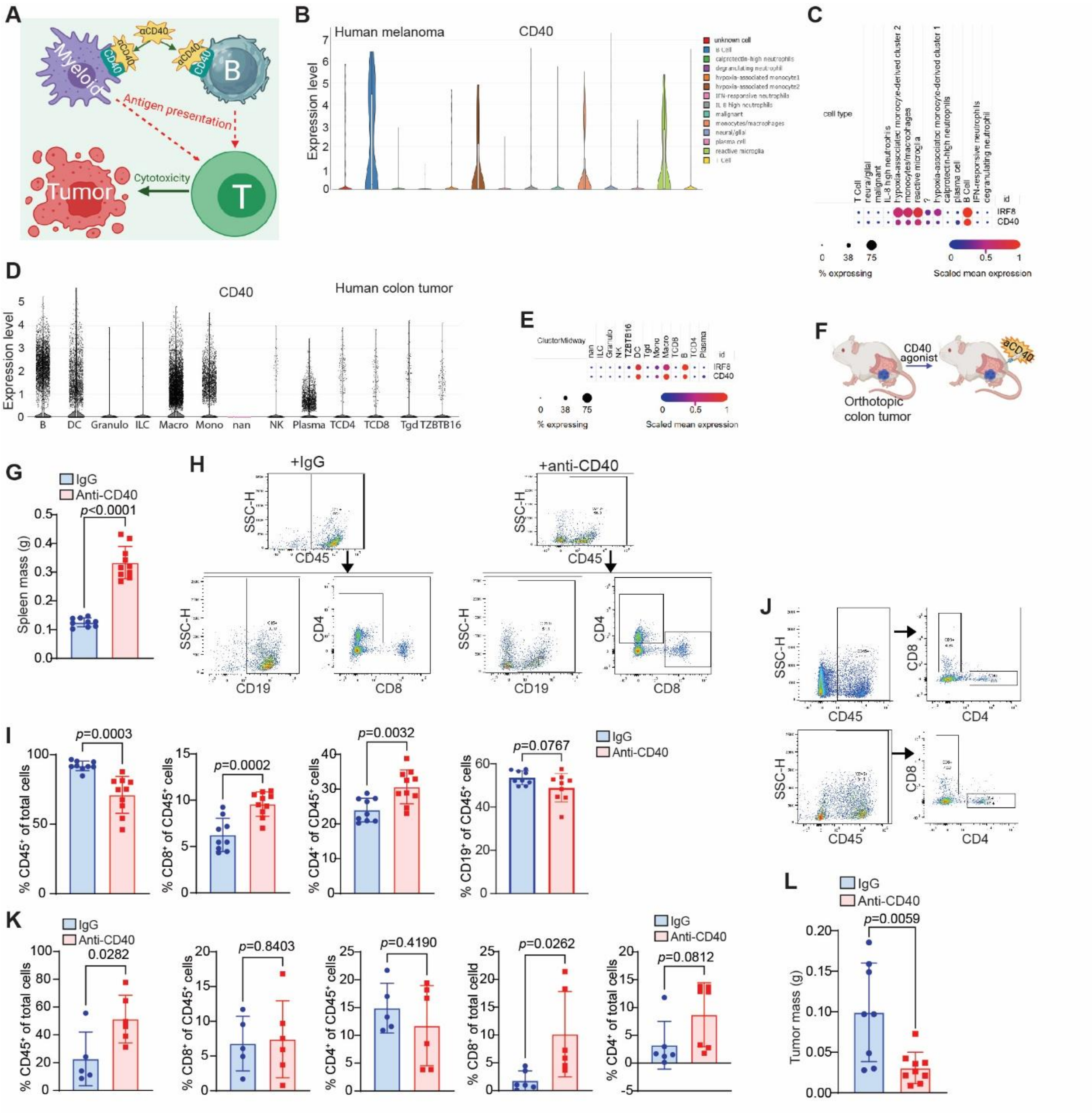

**Figure S7. CD40-mediated B cell activation restores anti-tumor immunity in tumors**

**A.** Schematic illustrating the antigen-presenting function of B cells and their capacity to activate T cell responses through antigen presentation and co-stimulatory signaling.

**B.** scRNA-seq feature plot showing CD40 expression in immune populations from human melanoma tumors. scRNA-seq data set was extracted from GEO data base (GEO accession #....)

**C.** Integrated scRNA-seq feature plots showing co-expression patterns of IRF8 and CD40 in human melanoma tumors. scRNA-seq data set was extracted from GEO data base (GEO accession #....)

**D.** scRNA-seq feature plot showing CD40 expression in immune populations from human colon tumors. scRNA-seq dataset was exctracted from GEO data base (GEO accession # ).

**E.** Integrated scRNA-seq feature plots showing IRF8 and CD40 expression in human colon tumors. scRNA-seq dataset was exctracted from GEO data base (GEO accession # ).

**F.** Experimental schema of the orthotopic colon tumor model. Colon tumor cells were surgically implanted into the mouse cecum, followed by treatment with agonistic anti-CD40 antibody or isotype control every 3 days.

**G.** Spleen masses from tumor-bearing mice treated with anti-CD40 or isotype control.

**H.** Representative flow cytometry gating strategy for analysis of splenic lymphocyte populations in mice treated with anti-CD40 or isotype control. Cells were stained with antibodies against CD45, CD19, CD4, and CD8.

**I.** Quantification of $CD45^+$ leukocytes, $CD19^+$ B cells, $CD4^+$ T cells, and $CD8^+$ T cells in spleens from anti-CD40-treated and control mice.

**J.** Representative flow cytometry gating strategy for analysis of tumor-infiltrating lymphocytes in mice treated with anti-CD40 or isotype control.

**K.** Quantification of tumor-infiltrating $CD45^+$ leukocytes, $CD4^+$ T cells, and $CD8^+$ T cells in anti-CD40-treated and control tumors.

**L.** Tumor masses from orthotopic colon tumor-bearing mice treated with anti-CD40 or IgG control.

**Table S1. IRF8-like motifs in tumor-infiltrating  myeloid and B cells**

| IRF8-like motif | Motif logo |
|---|---|
| MA0051.1 | 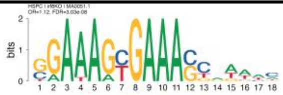 |
| MA0652.1 | 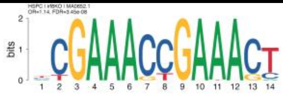 |
| MA0653.1 | 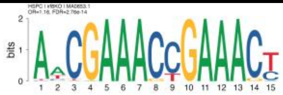 |
| MA0098.3 | 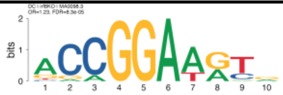 |
| MA0772.1 | 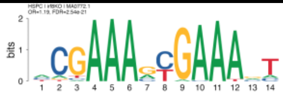 |
| MA1418.1 | 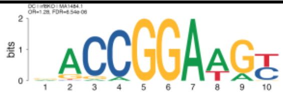 |
| MA1419.1 | 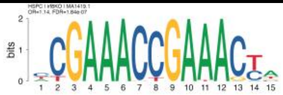 |
| MA1420.1 | 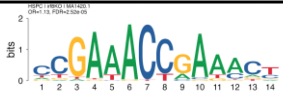 |
| MA1484.1 | 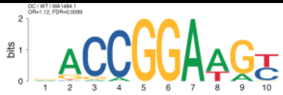 |
| MA1509.1 | 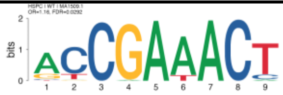 |
| AP-1(bZIP)/ThioMac-PU.1-ChIP-Seq(GSE21512)/Homer | 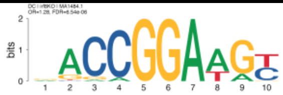 |
| bZIP:IRF(bZIP,IRF)/Th17-BatF-ChIP-Seq(GSE39756)/Homer | 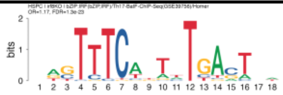 |
| E2A(bHLH),near_PU.1/Bcell-PU.1-ChIP-Seq(GSE21512)/Homer | 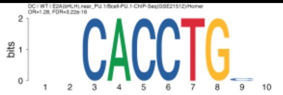 |
| EHF(ETS)/LoVo-EHF-ChIP-Seq(GSE49402)/Homer | 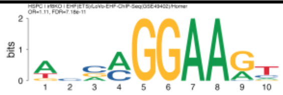 |
| ELF1(ETS)/Jurkat-ELF1-ChIP-Seq(SRA014231)/Homer | 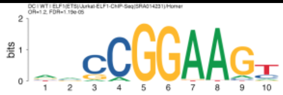 |
| ELF5(ETS)/T47D-ELF5-ChIP-Seq(GSE30407)/Homer | 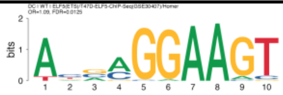 |
| Elk1(ETS)/Hela-Elk1-ChIP-Seq(GSE31477)/Homer | 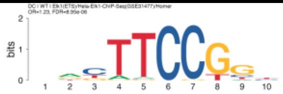 |

| | |
|---|---|
| Elk4(ETS)/Hela-Elk4-ChIP-Seq(GSE31477)/Homer | 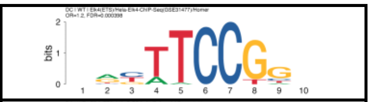 |
| ERG(ETS)/VCaP-ERG-ChIP-Seq(GSE14097)/Homer | 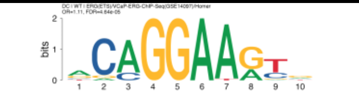 |
| ETS1(ETS)/Jurkat-ETS1-ChIP-Seq(GSE17954)/Homer | 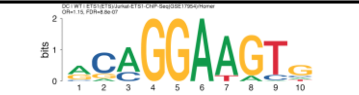 |
| Ets1-distal(ETS)/CD4+-PolII-ChIP-Seq(Barski et al.)/Homer | 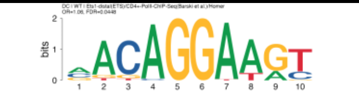 |
| ETS:E-box(ETS,bHLH)/HPC7-Scl-ChIP-Seq(GSE22178)/Homer | 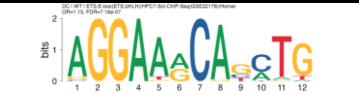 |
| ETS(ETS)/Promoter/Homer | 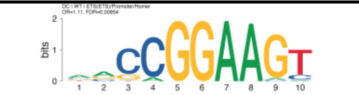 |
| ETS:RUNX(ETS,Runt)/Jurkat-RUNX1-ChIP-Seq(GSE17954)/Homer | 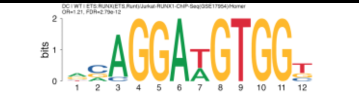 |
| ETV1(ETS)/GIST48-ETV1-ChIP-Seq(GSE22441)/Homer | 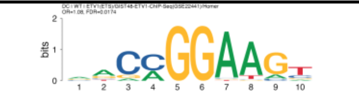 |
| EWS:ERG-fusion(ETS)/CADO_ES1-EWS:ERG-ChIP-Seq(SRA014231)/Homer | 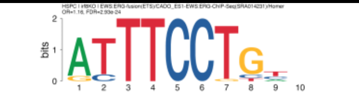 |
| EWS:FLI1-fusion(ETS)/SK_N_MC-EWS:FLI1-ChIP-Seq(SRA014231)/Homer | 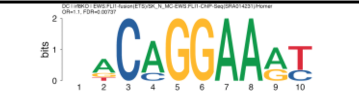 |
| Fli1(ETS)/CD8-FLI-ChIP-Seq(GSE20898)/Homer | 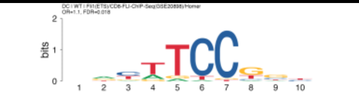 |
| GABPA(ETS)/Jurkat-GABPa-ChIP-Seq(GSE17954)/Homer | 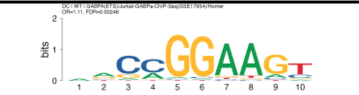 |
| IRF1(IRF)/PBMC-IRF1-ChIP-Seq(GSE43036)/Homer | 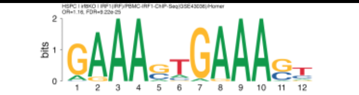 |
| IRF2(IRF)/Erythroblas-IRF2-ChIP-Seq(GSE36985)/Homer | 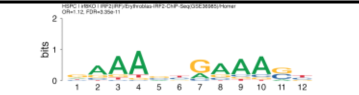 |
| IRF4(IRF)/GM12878-IRF4-ChIP-Seq(GSE32465)/Homer | 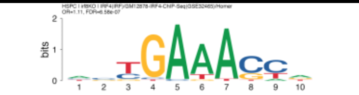 |
| ISRE(IRF)/ThioMac-LPS-Expression(GSE23622)/Homer | 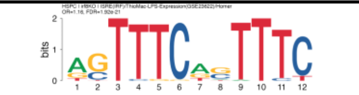 |
| PU.1-IRF(ETS:IRF)/Bcell-PU.1-ChIP-Seq(GSE21512)/Homer | 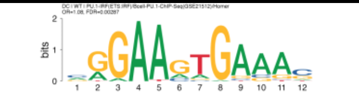 |
| PU.1(ETS)/ThioMac-PU.1-ChIP-Seq(GSE21512)/Homer | 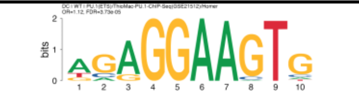 |
| SPDEF(ETS)/VCaP-SPDEF-ChIP-Seq(SRA014231)/Homer | 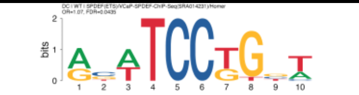 |

| | |
|---|---|
| T1ISRE(IRF)/ThioMac-Ifnb-Expression/Homer | 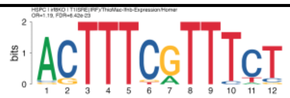 |
| TEAD(TEA)/Fibroblast-PU.1-ChIP-Seq(Unpublished)/Homer | 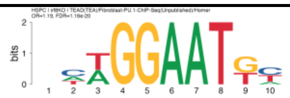 |

**Table S2. IRF8 motif enrichment in WT DC**

| Motif | motif DNA sequence logo | a | b | c | d | p-value | odds | FDR |
|---|---|---|---|---|---|---|---|---|
| MA0051.1 | | 1030 | 11939 | 10885 | 128143 | 0.64469505 | 1.01563171 | 0.69995463 |
| MA0652.1 | | 645 | 12324 | 7251 | 131777 | 0.23839731 | 0.95114801 | 0.31238269 |
| MA0653.1 | | 966 | 12003 | 10972 | 128056 | 0.07316326 | 0.93928464 | 0.10297052 |
| MA0098.3 | | 773 | 12196 | 7708 | 131320 | 0.04996237 | 1.0798467 | 0.07302193 |
| MA0772.1 | | 1099 | 11870 | 13659 | 125369 | 4.63E-07 | 0.84980408 | 2.51E-06 |
| MA1418.1 | | 1490 | 11479 | 16075 | 122953 | 0.81828018 | 0.99281943 | 0.86374018 |
| MA1419.1 | | 573 | 12396 | 6555 | 132473 | 0.12851202 | 0.93416809 | 0.17440917 |
| MA1420.1 | | 493 | 12476 | 5254 | 133774 | 0.88519484 | 1.00612813 | 0.89771479 |
| MA1484.1 | 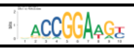 | 725 | 12244 | 6970 | 132058 | 0.00468921 | 1.1218551 | 0.00989945 |
| MA1509.1 | | 142 | 12827 | 1393 | 137635 | 0.31225886 | 1.09377758 | 0.38276893 |
| AP-1(bZIP)/ThioMac-PU.1-ChIP-Seq(GSE21512)/Homer | | 825 | 12144 | 15618 | 123410 | 3.50E-74 | 0.53681163 | 1.33E-72 |
| bZIP:IRF(bZIP,IRF)/Th17-BatF-ChIP-Seq(GSE39756)/Homer | | 1478 | 11491 | 18767 | 120261 | 7.19E-12 | 0.8242285 | 6.83E-11 |
| E2A(bHLH),near_PU.1/Bcell-PU.1-ChIP-Seq(GSE21512)/Homer | 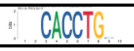 | 1573 | 11396 | 13522 | 125506 | 1.69E-17 | 1.28114607 | 3.22E-16 |
| EHF(ETS)/LoVo-EHF-ChIP-Seq(GSE49402)/Homer | | 1695 | 11274 | 17701 | 121327 | 0.27101425 | 1.03051111 | 0.34328472 |
| ELF1(ETS)/Jurkat-ELF1-ChIP-Seq(SRA014231)/Homer | 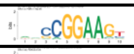 | 766 | 12203 | 6886 | 132142 | 3.43E-06 | 1.2045787 | 1.19E-05 |
| ELF5(ETS)/T47D-ELF5-ChIP-Seq(GSE30407)/Homer | 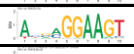 | 1246 | 11723 | 12354 | 126674 | 0.00624936 | 1.08985835 | 0.01249872 |
| Elk1(ETS)/Hela-Elk1-ChIP-Seq(GSE31477)/Homer | 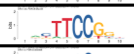 | 621 | 12348 | 5443 | 133585 | 2.12E-06 | 1.23427909 | 8.95E-06 |
| Elk4(ETS)/Hela-Elk4-ChIP-Seq(GSE31477)/Homer | 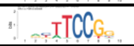 | 532 | 12437 | 4794 | 134234 | 0.00014669 | 1.19773202 | 0.00039815 |
| ERG(ETS)/VCaP-ERG-ChIP-Seq(GSE14097)/Homer | 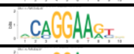 | 2270 | 10699 | 22289 | 116739 | 1.59E-05 | 1.111263 | 4.64E-05 |
| ETS1(ETS)/Jurkat-ETS1-ChIP-Seq(GSE17954)/Homer | 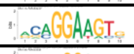 | 1984 | 10985 | 18921 | 120107 | 1.39E-07 | 1.14651673 | 8.80E-07 |
| Ets1-distal(ETS)/CD4+-PolII-ChIP-Seq(Barski et al.)/Homer | 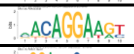 | 2099 | 10870 | 21486 | 117542 | 0.02918502 | 1.05638852 | 0.04480708 |
| ETS:E-box(ETS,bHLH)/HPC7-Scl-ChIP-Seq(GSE22178)/Homer | 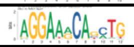 | 1896 | 11073 | 17996 | 121032 | 9.42E-08 | 1.15154929 | 7.16E-07 |
| ETS(ETS)/Promoter/Homer | 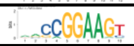 | 940 | 12029 | 9158 | 129870 | 0.00426645 | 1.10813585 | 0.00953676 |
| ETS:RUNX(ETS,Runt)/Jurkat-RUNX1-ChIP-Seq(GSE17954)/Homer | 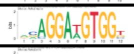 | 1944 | 11025 | 17638 | 121390 | 2.20E-13 | 1.21353005 | 2.79E-12 |
| ETV1(ETS)/GIST48-ETV1-ChIP-Seq(GSE22441)/Homer | 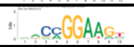 | 1527 | 11442 | 15323 | 123705 | 0.00964168 | 1.07742757 | 0.01744686 |
| EWS:ERG-fusion(ETS)/CADO_ES1-EWS:ERG-ChIP-Seq(SRA014231)/Homer | | 1981 | 10988 | 21570 | 117458 | 0.47742098 | 0.98174134 | 0.53490408 |
| EWS:FLI1-fusion(ETS)/SK_N_MC-EWS:FLI1-ChIP-Seq(SRA014231)/Homer | | 1836 | 11133 | 19258 | 119770 | 0.33902036 | 1.02564774 | 0.40258668 |
| Fli1(ETS)/CD8-FLI-ChIP-Seq(GSE20898)/Homer | 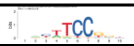 | 825 | 12144 | 8070 | 130958 | 0.01039702 | 1.10239385 | 0.01795849 |
| GABPA(ETS)/Jurkat-GABPa-ChIP-Seq(GSE17954)/Homer | 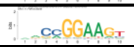 | 1263 | 11706 | 12326 | 126702 | 0.0009709 | 1.10909909 | 0.00245961 |
| IRF1(IRF)/PBMC-IRF1-ChIP-Seq(GSE43036)/Homer | | 2134 | 10835 | 24160 | 114868 | 0.00785539 | 0.93643417 | 0.01492525 |
| IRF2(IRF)/Erythroblas-IRF2-ChIP-Seq(GSE36985)/Homer | | 1522 | 11447 | 16261 | 122767 | 0.89771479 | 1.00382355 | 0.89771479 |

| | | | | | | | | |
|---|---|---|---|---|---|---|---|---|
| IRF4(IRF)/GM12878-IRF4-ChIP-Seq(GSE32465)/Homer | | 821 | 12148 | 9032 | 129996 | 0.47859839 | 0.97271094 | 0.53490408 |
| ISRE(IRF)/ThioMac-LPS-Expression(GSE23622)/Homer | | 1619 | 11350 | 18294 | 120734 | 0.02947834 | 0.94137522 | 0.04480708 |
| PU.1-IRF(ETS:IRF)/Bcell-PU.1-ChIP-Seq(GSE21512)/Homer | 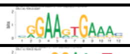 | 2320 | 10649 | 23314 | 115714 | 0.00120827 | 1.08131842 | 0.00286963 |
| PU.1(ETS)/ThioMac-PU.1-ChIP-Seq(GSE21512)/Homer | 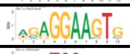 | 2065 | 10904 | 20138 | 118890 | 1.18E-05 | 1.11808248 | 3.73E-05 |
| SPDEF(ETS)/VCaP-SPDEF-ChIP-Seq(SRA014231)/Homer | 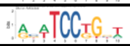 | 1367 | 11602 | 13797 | 125231 | 0.02632412 | 1.06947274 | 0.04349203 |
| T1ISRE(IRF)/ThioMac-Ifnb-Expression/Homer | | 1214 | 11755 | 14813 | 124215 | 3.32E-06 | 0.86601934 | 1.19E-05 |
| TEAD(TEA)/Fibroblast-PU.1-ChIP-Seq(Unpublished)/Homer | | 1033 | 11936 | 12857 | 126171 | 8.46E-07 | 0.84930167 | 4.02E-06 |

a: Number of target peaks containing the motif

b: Number of target peaks not containing the motif

c: Number of background peaks containing the motif

d: Number of background peaks not containing the motif

| Table S3. IRF8 motif enrichment in WT IMCs | | | | | | | | |
|---|---|---|---|---|---|---|---|---|
| Motif | motif DNA sequence logo | a | b | c | d | pvalue | odds | FDR |
| MA0051.1 | 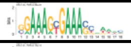 | 3132 | 31234 | 8783 | 108848 | 8.44E-23 | 1.24272321 | 4.58E-22 |
| MA0652.1 | 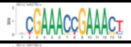 | 2046 | 32320 | 5850 | 111781 | 1.32E-12 | 1.20963497 | 4.15E-12 |
| MA0653.1 | 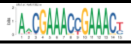 | 3089 | 31277 | 8849 | 108782 | 1.99E-18 | 1.2141276 | 7.55E-18 |
| MA0098.3 | | 1860 | 32506 | 6621 | 111010 | 0.12780934 | 0.9593488 | 0.1674743 |
| MA0772.1 | 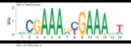 | 3989 | 30377 | 10769 | 106862 | 4.01E-40 | 1.30306539 | 5.07E-39 |
| MA1418.1 | 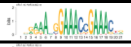 | 4438 | 29928 | 13127 | 104504 | 9.08E-19 | 1.18048991 | 3.83E-18 |
| MA1419.1 | 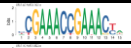 | 1860 | 32506 | 5268 | 112363 | 1.42E-12 | 1.22048429 | 4.15E-12 |
| MA1420.1 | 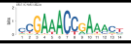 | 1440 | 32926 | 4307 | 113324 | 7.82E-06 | 1.15069039 | 1.86E-05 |
| MA1484.1 | | 1676 | 32690 | 6019 | 111612 | 0.07571085 | 0.95073018 | 0.11987551 |
| MA1509.1 | 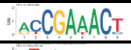 | 387 | 33979 | 1148 | 116483 | 0.01538953 | 1.15566298 | 0.02924012 |
| AP-1(bZIP)/ThioMac-PU.1-ChIP-Seq(GSE21512)/Homer | 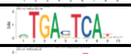 | 5303 | 29063 | 11140 | 106491 | 7.21E-200 | 1.74419658 | 2.74E-198 |
| bZIP:IRF(bZIP,IRF)/Th17-BatF-ChIP-Seq(GSE39756)/Homer | 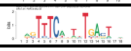 | 5356 | 29010 | 14889 | 102742 | 1.55E-43 | 1.27402124 | 2.95E-42 |
| E2A(bHLH),near_PU.1/Bcell-PU.1-ChIP-Seq(GSE21512)/Homer | | 3473 | 30893 | 11622 | 106009 | 0.21864682 | 1.02543223 | 0.25964309 |
| EHF(ETS)/LoVo-EHF-ChIP-Seq(GSE49402)/Homer | | 4494 | 29872 | 14902 | 102729 | 0.04615348 | 1.03709375 | 0.07625357 |
| ELF1(ETS)/Jurkat-ELF1-ChIP-Seq(SRA014231)/Homer | | 1678 | 32688 | 5974 | 111657 | 0.14478526 | 0.95943019 | 0.18339466 |
| ELF5(ETS)/T47D-ELF5-ChIP-Seq(GSE30407)/Homer | | 3121 | 31245 | 10479 | 107152 | 0.32304988 | 1.02139557 | 0.35073987 |
| Elk1(ETS)/Hela-Elk1-ChIP-Seq(GSE31477)/Homer | | 1317 | 33049 | 4747 | 112884 | 0.0906883 | 0.9476576 | 0.1324091 |
| Elk4(ETS)/Hela-Elk4-ChIP-Seq(GSE31477)/Homer | | 1140 | 33226 | 4186 | 113445 | 0.03281274 | 0.92985077 | 0.05667655 |
| ERG(ETS)/VCaP-ERG-ChIP-Seq(GSE14097)/Homer | | 5450 | 28916 | 19109 | 98522 | 0.08770368 | 0.97174733 | 0.1324091 |
| ETS1(ETS)/Jurkat-ETS1-ChIP-Seq(GSE17954)/Homer | | 4479 | 29887 | 16426 | 101205 | 9.66E-06 | 0.92335697 | 2.16E-05 |
| Ets1-distal(ETS)/CD4+-PolII-ChIP-Seq(Barski et al.)/Homer | | 5238 | 29128 | 18347 | 99284 | 0.11138841 | 0.97312556 | 0.15116999 |
| ETS:E-box(ETS,bHLH)/HPC7-Scl-ChIP-Seq(GSE22178)/Homer | | 4505 | 29861 | 15387 | 102244 | 0.89153774 | 1.00247678 | 0.89153774 |
| ETS(ETS)/Promoter/Homer | | 2215 | 32151 | 7883 | 109748 | 0.09408015 | 0.95911795 | 0.1324091 |
| ETS:RUNX(ETS,Runt)/Jurkat-RUNX1-ChIP-Seq(GSE17954)/Homer | | 4353 | 30013 | 15229 | 102402 | 0.17559174 | 0.97523591 | 0.21524148 |
| ETV1(ETS)/GIST48-ETV1-ChIP-Seq(GSE22441)/Homer | | 3692 | 30674 | 13158 | 104473 | 0.0217356 | 0.95568338 | 0.03933108 |
| EWS:ERG-fusion(ETS)/CADO_ES1-EWS:ERG-ChIP-Seq(SRA014231)/Homer | 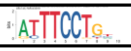 | 5498 | 28868 | 18053 | 99578 | 0.00346406 | 1.05051739 | 0.00731302 |
| EWS:FLI1-fusion(ETS)/SK_N_MC-EWS:FLI1-ChIP-Seq(SRA014231)/Homer | | 4730 | 29636 | 16364 | 101267 | 0.48911514 | 0.98768466 | 0.50233447 |
| Fli1(ETS)/CD8-FLI-ChIP-Seq(GSE20898)/Homer | | 1907 | 32459 | 6988 | 110643 | 0.00658598 | 0.93022278 | 0.01317196 |
| GABPA(ETS)/Jurkat-GABPa-ChIP-Seq(GSE17954)/Homer | | 3017 | 31349 | 10572 | 107059 | 0.2372063 | 0.97456805 | 0.27314665 |
| IRF1(IRF)/PBMC-IRF1-ChIP-Seq(GSE43036)/Homer | 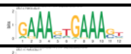 | 6677 | 27689 | 19617 | 98014 | 8.44E-32 | 1.20479556 | 6.42E-31 |
| IRF2(IRF)/Erythroblas-IRF2-ChIP-Seq(GSE36985)/Homer | 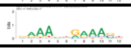 | 4474 | 29892 | 13309 | 104322 | 1.23E-17 | 1.17316224 | 4.24E-17 |

| | | | | | | | | |
|---|---|---|---|---|---|---|---|---|
| IRF4(IRF)/GM12878-IRF4-ChIP-Seq(GSE32465)/Homer | 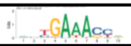 | 2456 | 31910 | 7397 | 110234 | 1.93E-08 | 1.14696688 | 5.25E-08 |
| ISRE(IRF)/ThioMac-LPS-Expression(GSE23622)/Homer | 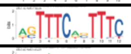 | 5030 | 29336 | 14883 | 102748 | 2.51E-21 | 1.18368276 | 1.19E-20 |
| PU.1-IRF(ETS:IRF)/Bcell-PU.1-ChIP-Seq(GSE21512)/Homer | 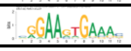 | 6130 | 28236 | 19504 | 98127 | 5.17E-08 | 1.09224365 | 1.31E-07 |
| PU.1(ETS)/ThioMac-PU.1-ChIP-Seq(GSE21512)/Homer | | 4968 | 29398 | 17235 | 100396 | 0.37125107 | 0.98439386 | 0.39187613 |
| SPDEF(ETS)/VCaP-SPDEF-ChIP-Seq(SRA014231)/Homer | | 3485 | 30881 | 11679 | 105952 | 0.24767599 | 1.02380035 | 0.27681434 |
| T1ISRE(IRF)/ThioMac-Ifnb-Expression/Homer | 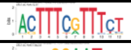 | 4230 | 30136 | 11797 | 105834 | 1.05E-32 | 1.25924423 | 1.00E-31 |
| TEAD(TEA)/Fibroblast-PU.1-ChIP-Seq(Unpublished)/Homer | 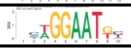 | 3616 | 30750 | 10274 | 107357 | 2.15E-23 | 1.22879858 | 1.36E-22 |

| Table S4. IRF8 motif enrichment in WT plasmblasts | | | | | | | | |
|---|---|---|---|---|---|---|---|---|
| Motif | Motif DNA sequence logo | a | b | c | d | p-value | odds | FDR |
| MA0051.1 | | 1716 | 20052 | 10199 | 120030 | 0.79578236 | 1.00714468 | 0.81262192 |
| MA0652.1 | 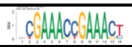 | 1210 | 20558 | 6686 | 123543 | 0.00958508 | 1.08758839 | 0.01456932 |
| MA0653.1 | | 1734 | 20034 | 10204 | 120025 | 0.50488223 | 1.01808288 | 0.54815785 |
| MA0098.3 | 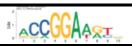 | 1348 | 20420 | 7133 | 123096 | 2.71E-05 | 1.13917562 | 5.42E-05 |
| MA0772.1 | | 2064 | 19704 | 12694 | 117535 | 0.22557711 | 0.96988622 | 0.25969554 |
| MA1418.1 | 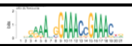 | 2625 | 19143 | 14940 | 115289 | 0.01253826 | 1.05818083 | 0.01832515 |
| MA1419.1 | | 1071 | 20697 | 6057 | 124172 | 0.08330842 | 1.06084748 | 0.105524 |
| MA1420.1 | | 864 | 20904 | 4883 | 125346 | 0.11551583 | 1.06099436 | 0.14160005 |
| MA1484.1 | 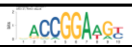 | 1284 | 20484 | 6411 | 123818 | 2.69E-09 | 1.21059373 | 1.28E-08 |
| MA1509.1 | | 226 | 21542 | 1309 | 128920 | 0.66030471 | 1.03324898 | 0.69698831 |
| AP-1(bZIP)/ThioMac-PU.1-ChIP-Seq(GSE21512)/Homer | | 1689 | 20079 | 14754 | 115475 | 9.15E-60 | 0.65837492 | 3.48E-58 |
| bZIP:IRF(bZIP,IRF)/Th17-BatF-ChIP-Seq(GSE39756)/Homer | | 2888 | 18880 | 17357 | 112872 | 0.81262192 | 0.99473323 | 0.81262192 |
| E2A(bHLH),near_PU.1/Bcell-PU.1-ChIP-Seq(GSE21512)/Homer | 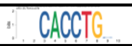 | 2248 | 19520 | 12847 | 117382 | 0.03524438 | 1.05225073 | 0.04783166 |
| EHF(ETS)/LoVo-EHF-ChIP-Seq(GSE49402)/Homer | 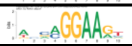 | 3032 | 18736 | 16364 | 113865 | 3.38E-08 | 1.12606343 | 1.28E-07 |
| ELF1(ETS)/Jurkat-ELF1-ChIP-Seq(SRA014231)/Homer | 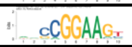 | 1225 | 20543 | 6427 | 123802 | 1.95E-05 | 1.14862334 | 4.63E-05 |
| ELF5(ETS)/T47D-ELF5-ChIP-Seq(GSE30407)/Homer | 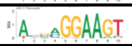 | 2112 | 19656 | 11488 | 118741 | 3.05E-05 | 1.11061836 | 5.80E-05 |
| Elk1(ETS)/Hela-Elk1-ChIP-Seq(GSE31477)/Homer | 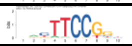 | 983 | 20785 | 5081 | 125148 | 2.54E-05 | 1.1648405 | 5.37E-05 |
| Elk4(ETS)/Hela-Elk4-ChIP-Seq(GSE31477)/Homer | 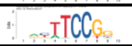 | 864 | 20904 | 4462 | 125767 | 7.37E-05 | 1.16495764 | 0.00012732 |
| ERG(ETS)/VCaP-ERG-ChIP-Seq(GSE14097)/Homer | 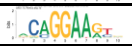 | 3847 | 17921 | 20712 | 109517 | 8.19E-11 | 1.13503121 | 4.45E-10 |
| ETS1(ETS)/Jurkat-ETS1-ChIP-Seq(GSE17954)/Homer | 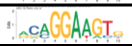 | 3376 | 18392 | 17529 | 112700 | 1.16E-15 | 1.18017082 | 1.47E-14 |
| Ets1-distal(ETS)/CD4+-PolII-ChIP-Seq(Barski et al.)/Homer | 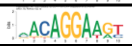 | 3743 | 18025 | 19842 | 110387 | 2.81E-13 | 1.15528916 | 2.14E-12 |
| ETS:E-box(ETS,bHLH)/HPC7-Scl-ChIP-Seq(GSE22178)/Homer | 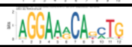 | 3045 | 18723 | 16847 | 113382 | 2.41E-05 | 1.094561 | 5.37E-05 |
| ETS(ETS)/Promoter/Homer | 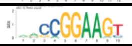 | 1620 | 20148 | 8478 | 121751 | 4.56E-07 | 1.15464282 | 1.44E-06 |
| ETS:RUNX(ETS,Runt)/Jurkat-RUNX1-ChIP-Seq(GSE17954)/Homer | 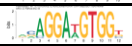 | 3196 | 18572 | 16386 | 113843 | 3.46E-17 | 1.1955862 | 6.57E-16 |
| ETV1(ETS)/GIST48-ETV1-ChIP-Seq(GSE22441)/Homer | 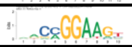 | 2711 | 19057 | 14139 | 116090 | 7.49E-12 | 1.16806073 | 4.74E-11 |
| EWS:ERG-fusion(ETS)/CADO_ES1-EWS:ERG-ChIP-Seq(SRA014231)/Homer | 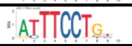 | 3600 | 18168 | 19951 | 110278 | 5.02E-06 | 1.09528061 | 1.41E-05 |
| EWS:FLI1-fusion(ETS)/SK_N_MC-EWS:FLI1-ChIP-Seq(SRA014231)/Homer | 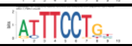 | 3234 | 18534 | 17860 | 112369 | 7.46E-06 | 1.0978489 | 1.89E-05 |
| Fli1(ETS)/CD8-FLI-ChIP-Seq(GSE20898)/Homer | 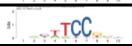 | 1422 | 20346 | 7473 | 122756 | 5.21E-06 | 1.14803135 | 1.41E-05 |
| GABPA(ETS)/Jurkat-GABPa-ChIP-Seq(GSE17954)/Homer | 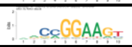 | 2180 | 19588 | 11409 | 118820 | 3.27E-09 | 1.15902447 | 1.38E-08 |
| IRF1(IRF)/PBMC-IRF1-ChIP-Seq(GSE43036)/Homer | 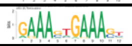 | 3909 | 17859 | 22385 | 107844 | 0.00580113 | 1.05450718 | 0.00918512 |
| IRF2(IRF)/Erythroblas-IRF2-ChIP-Seq(GSE36985)/Homer | 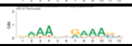 | 2683 | 19085 | 15100 | 115129 | 0.00202126 | 1.07186706 | 0.00333947 |

| | | | | | | | | |
|---|---|---|---|---|---|---|---|---|
| IRF4(IRF)/GM12878-IRF4-ChIP-Seq(GSE32465)/Homer | 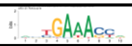 | 1493 | 20275 | 8360 | 121869 | 0.01535456 | 1.07347498 | 0.02161012 |
| ISRE(IRF)/ThioMac-LPS-Expression(GSE23622)/Homer | | 2943 | 18825 | 16970 | 113259 | 0.04828382 | 1.04339323 | 0.06326846 |
| PU.1-IRF(ETS:IRF)/Bcell-PU.1-ChIP-Seq(GSE21512)/Homer | 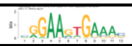 | 3941 | 17827 | 21693 | 108536 | 1.68E-07 | 1.10608503 | 5.79E-07 |
| PU.1(ETS)/ThioMac-PU.1-ChIP-Seq(GSE21512)/Homer | 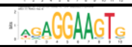 | 3554 | 18214 | 18649 | 111580 | 1.81E-14 | 1.16748748 | 1.72E-13 |
| SPDEF(ETS)/VCaP-SPDEF-ChIP-Seq(SRA014231)/Homer | 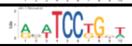 | 2336 | 19432 | 12828 | 117401 | 6.80E-05 | 1.10021201 | 0.00012309 |
| T1ISRE(IRF)/ThioMac-Ifnb-Expression/Homer | | 2244 | 19524 | 13783 | 116446 | 0.22401522 | 0.97102769 | 0.25969554 |
| TEAD(TEA)/Fibroblast-PU.1-ChIP-Seq(Unpublished)/Homer | | 1942 | 19826 | 11948 | 118281 | 0.23235917 | 0.96968533 | 0.25969554 |

| **Table S5. IRF8 motif enrichment in IRF8 KO DCs** | | | | | | | | |
|---|---|---|---|---|---|---|---|---|
| Motif | motif DNA sequence logo | a | b | c | d | pvalue | odds | FDR |
| MA0051.1 | | 472 | 6452 | 11443 | 133630 | 0.00106217 | 0.85428164 | 0.00167076 |
| MA0652.1 | | 281 | 6643 | 7615 | 137458 | 7.01E-06 | 0.76353685 | 1.90E-05 |
| MA0653.1 | | 462 | 6462 | 11476 | 133597 | 0.00014635 | 0.83227567 | 0.00025278 |
| MA0098.3 | 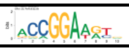 | 465 | 6459 | 8016 | 137057 | 4.07E-05 | 1.23092088 | 8.30E-05 |
| MA0772.1 | | 544 | 6380 | 14214 | 130859 | 4.57E-08 | 0.78499386 | 2.17E-07 |
| MA1418.1 | 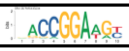 | 724 | 6200 | 16841 | 128232 | 0.003238 | 0.88918182 | 0.00456455 |
| MA1419.1 | | 263 | 6661 | 6865 | 138208 | 0.00024437 | 0.79487908 | 0.00040374 |
| MA1420.1 | | 235 | 6689 | 5512 | 139561 | 0.08733989 | 0.88952852 | 0.1005732 |
| MA1484.1 | | 438 | 6486 | 7257 | 137816 | 2.06E-06 | 1.28244417 | 6.54E-06 |
| MA1509.1 | | 81 | 6843 | 1454 | 143619 | 0.1755655 | 1.16918987 | 0.19061397 |
| AP-1(bZIP)/ThioMac-PU.1-ChIP-Seq(GSE21512)/Homer | 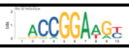 | 473 | 6451 | 15970 | 129103 | 4.95E-31 | 0.59274459 | 1.88E-29 |
| bZIP:IRF(bZIP,IRF)/Th17-BatF-ChIP-Seq(GSE39756)/Homer | | 818 | 6106 | 19427 | 125646 | 0.00013336 | 0.86644404 | 0.00024132 |
| E2A(bHLH),near_PU.1/Bcell-PU.1-ChIP-Seq(GSE21512)/Homer | 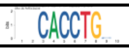 | 845 | 6079 | 14250 | 130823 | 3.40E-10 | 1.27612279 | 3.23E-09 |
| EHF(ETS)/LoVo-EHF-ChIP-Seq(GSE49402)/Homer | | 934 | 5990 | 18462 | 126611 | 0.06524078 | 1.06934895 | 0.07747343 |
| ELF1(ETS)/Jurkat-ELF1-ChIP-Seq(SRA014231)/Homer | 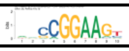 | 487 | 6437 | 7165 | 137908 | 1.46E-13 | 1.4561761 | 2.77E-12 |
| ELF5(ETS)/T47D-ELF5-ChIP-Seq(GSE30407)/Homer | | 654 | 6270 | 12946 | 132127 | 0.13706826 | 1.06456974 | 0.15319394 |
| Elk1(ETS)/Hela-Elk1-ChIP-Seq(GSE31477)/Homer | 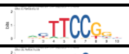 | 398 | 6526 | 5666 | 139407 | 5.14E-13 | 1.50050805 | 6.51E-12 |
| Elk4(ETS)/Hela-Elk4-ChIP-Seq(GSE31477)/Homer | 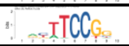 | 340 | 6584 | 4986 | 140087 | 6.72E-10 | 1.45087215 | 5.11E-09 |
| ERG(ETS)/VCaP-ERG-ChIP-Seq(GSE14097)/Homer | 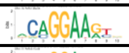 | 1275 | 5649 | 23284 | 121789 | 2.85E-07 | 1.18052584 | 1.08E-06 |
| ETS1(ETS)/Jurkat-ETS1-ChIP-Seq(GSE17954)/Homer | 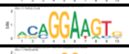 | 1085 | 5839 | 19820 | 125253 | 3.12E-06 | 1.17426231 | 9.11E-06 |
| Ets1-distal(ETS)/CD4+-PolII-ChIP-Seq(Barski et al.)/Homer | 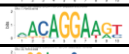 | 1157 | 5767 | 22428 | 122645 | 0.00533284 | 1.09711387 | 0.00723204 |
| ETS:E-box(ETS,bHLH)/HPC7-Scl-ChIP-Seq(GSE22178)/Homer | 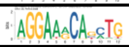 | 966 | 5958 | 18926 | 126147 | 0.02998192 | 1.08069462 | 0.03675203 |
| ETS(ETS)/Promoter/Homer | 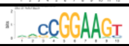 | 569 | 6355 | 9529 | 135544 | 1.82E-07 | 1.27358679 | 7.68E-07 |
| ETS:RUNX(ETS,Runt)/Jurkat-RUNX1-ChIP-Seq(GSE17954)/Homer | 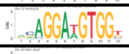 | 1006 | 5918 | 18576 | 126497 | 3.89E-05 | 1.15755124 | 8.30E-05 |
| ETV1(ETS)/GIST48-ETV1-ChIP-Seq(GSE22441)/Homer | 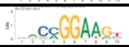 | 913 | 6011 | 15937 | 129136 | 2.58E-08 | 1.23072763 | 1.45E-07 |
| EWS:ERG-fusion(ETS)/CADO_ES1-EWS:ERG-ChIP-Seq(SRA014231)/Homer | | 1062 | 5862 | 22489 | 122584 | 0.72115735 | 0.9875111 | 0.72115735 |
| EWS:FLI1-fusion(ETS)/SK_N_MC-EWS:FLI1-ChIP-Seq(SRA014231)/Homer | 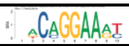 | 1039 | 5885 | 20055 | 125018 | 0.00581893 | 1.10060046 | 0.00737064 |
| Fli1(ETS)/CD8-FLI-ChIP-Seq(GSE20898)/Homer | 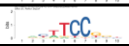 | 515 | 6409 | 8380 | 136693 | 2.67E-08 | 1.31074086 | 1.45E-07 |
| GABPA(ETS)/Jurkat-GABPa-ChIP-Seq(GSE17954)/Homer | 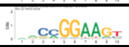 | 733 | 6191 | 12856 | 132217 | 1.67E-06 | 1.2176535 | 5.79E-06 |
| IRF1(IRF)/PBMC-IRF1-ChIP-Seq(GSE43036)/Homer | | 1073 | 5851 | 25221 | 119852 | 4.15E-05 | 0.87147163 | 8.30E-05 |
| IRF2(IRF)/Erythroblas-IRF2-ChIP-Seq(GSE36985)/Homer | | 738 | 6186 | 17045 | 128028 | 0.00551919 | 0.89606831 | 0.00723204 |
| IRF4(IRF)/GM12878-IRF4-ChIP-Seq(GSE32465)/Homer | | 430 | 6494 | 9423 | 135650 | 0.35538827 | 0.95320445 | 0.36499336 |

| | | | | | | | | |
|---|---|---|---|---|---|---|---|---|
| ISRE(IRF)/ThioMac-LPS-Expression(GSE23622)/Homer | | 818 | 6106 | 19095 | 125978 | 0.00109919 | 0.88385279 | 0.00167076 |
| PU.1-IRF(ETS:IRF)/Bcell-PU.1-ChIP-Seq(GSE21512)/Homer | | 1201 | 5723 | 24433 | 120640 | 0.27829307 | 1.03618039 | 0.2937538 |
| PU.1(ETS)/ThioMac-PU.1-ChIP-Seq(GSE21512)/Homer | | 1097 | 5827 | 21106 | 123967 | 0.00324323 | 1.10578995 | 0.00456455 |
| SPDEF(ETS)/VCaP-SPDEF-ChIP-Seq(SRA014231)/Homer | 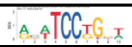 | 788 | 6136 | 14376 | 130697 | 8.80E-05 | 1.16751728 | 0.00016714 |
| T1ISRE(IRF)/ThioMac-Ifnb-Expression/Homer | | 629 | 6295 | 15398 | 129675 | 4.00E-05 | 0.84150091 | 8.30E-05 |
| TEAD(TEA)/Fibroblast-PU.1-ChIP-Seq(Unpublished)/Homer | | 534 | 6390 | 13356 | 131717 | 1.77E-05 | 0.82417019 | 4.48E-05 |

| Table S6. IRF8 motif enrichment in IRF8 KO IMCs | | | | | | | | |
|---|---|---|---|---|---|---|---|---|
| Motif | motif DNA sequence logo | a | b | c | d | pvalue | odds | FDR |
| MA0051.1 | 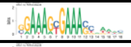 | 4633 | 50818 | 7282 | 89264 | 1.60E-08 | 1.11755456 | 3.03E-08 |
| MA0652.1 | 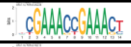 | 3116 | 52335 | 4780 | 91766 | 1.91E-08 | 1.14302724 | 3.45E-08 |
| MA0653.1 | 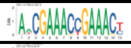 | 4748 | 50703 | 7190 | 89356 | 1.02E-14 | 1.16377355 | 2.76E-14 |
| MA0098.3 | 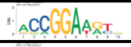 | 3235 | 52216 | 5246 | 91300 | 0.00110693 | 1.07823267 | 0.00150226 |
| MA0772.1 | 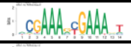 | 5923 | 49528 | 8835 | 87711 | 5.35E-22 | 1.18722754 | 2.54E-21 |
| MA1418.1 | 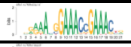 | 6872 | 48579 | 10693 | 85853 | 1.34E-14 | 1.13576432 | 3.40E-14 |
| MA1419.1 | 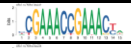 | 2812 | 52639 | 4316 | 92230 | 1.12E-07 | 1.14155271 | 1.84E-07 |
| MA1420.1 | 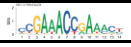 | 2251 | 53200 | 3496 | 93050 | 1.79E-05 | 1.12617757 | 2.52E-05 |
| MA1484.1 | | 2801 | 52650 | 4894 | 91652 | 0.88406218 | 0.99630593 | 0.90795575 |
| MA1509.1 | | 564 | 54887 | 971 | 95575 | 0.83123226 | 1.01143055 | 0.87741183 |
| AP-1(bZIP)/ThioMac-PU.1-ChIP-Seq(GSE21512)/Homer | 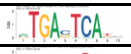 | 6904 | 48547 | 9539 | 87007 | 1.55E-53 | 1.29711663 | 5.91E-52 |
| bZIP:IRF(bZIP,IRF)/Th17-BatF-ChIP-Seq(GSE39756)/Homer | 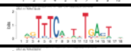 | 8042 | 47409 | 12203 | 84343 | 1.37E-24 | 1.17241655 | 1.30E-23 |
| E2A(bHLH),near_PU.1/Bcell-PU.1-ChIP-Seq(GSE21512)/Homer | 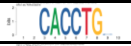 | 5684 | 49767 | 9411 | 87135 | 0.00166288 | 1.05747256 | 0.00217895 |
| EHF(ETS)/LoVo-EHF-ChIP-Seq(GSE49402)/Homer | 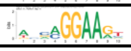 | 7493 | 47958 | 11903 | 84643 | 3.21E-11 | 1.11103522 | 7.18E-11 |
| ELF1(ETS)/Jurkat-ELF1-ChIP-Seq(SRA014231)/Homer | | 2741 | 52710 | 4911 | 91635 | 0.2184828 | 0.97030174 | 0.23943596 |
| ELF5(ETS)/T47D-ELF5-ChIP-Seq(GSE30407)/Homer | 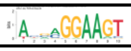 | 5204 | 50247 | 8396 | 88150 | 6.52E-06 | 1.08736704 | 9.53E-06 |
| Elk1(ETS)/Hela-Elk1-ChIP-Seq(GSE31477)/Homer | | 2167 | 53284 | 3897 | 92649 | 0.22053312 | 0.96687792 | 0.23943596 |
| Elk4(ETS)/Hela-Elk4-ChIP-Seq(GSE31477)/Homer | | 1939 | 53512 | 3387 | 93159 | 0.91921786 | 0.99663526 | 0.91921786 |
| ERG(ETS)/VCaP-ERG-ChIP-Seq(GSE14097)/Homer | 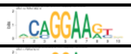 | 9500 | 45951 | 15059 | 81487 | 6.34E-15 | 1.11871424 | 1.85E-14 |
| ETS1(ETS)/Jurkat-ETS1-ChIP-Seq(GSE17954)/Homer | 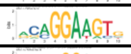 | 8043 | 47408 | 12862 | 83684 | 1.35E-10 | 1.10382215 | 2.70E-10 |
| Ets1-distal(ETS)/CD4+-PolII-ChIP-Seq(Barski et al.)/Homer | 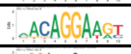 | 9200 | 46251 | 14385 | 82161 | 2.42E-18 | 1.13610977 | 9.18E-18 |
| ETS:E-box(ETS,bHLH)/HPC7-Scl-ChIP-Seq(GSE22178)/Homer | 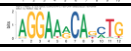 | 7775 | 47676 | 12117 | 84429 | 3.75E-16 | 1.13630481 | 1.19E-15 |
| ETS(ETS)/Promoter/Homer | | 3753 | 51698 | 6345 | 90201 | 0.13987417 | 1.03203579 | 0.16106723 |
| ETS:RUNX(ETS,Runt)/Jurkat-RUNX1-ChIP-Seq(GSE17954)/Homer | 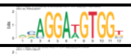 | 7326 | 48125 | 12256 | 84290 | 0.00389138 | 1.04692701 | 0.00477008 |
| ETV1(ETS)/GIST48-ETV1-ChIP-Seq(GSE22441)/Homer | 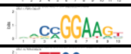 | 6466 | 48985 | 10384 | 86162 | 7.08E-08 | 1.09527363 | 1.22E-07 |
| EWS:ERG-fusion(ETS)/CADO_ES1-EWS:ERG-ChIP-Seq(SRA014231)/Homer | 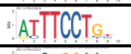 | 9302 | 46149 | 14249 | 82297 | 2.31E-25 | 1.16415359 | 2.93E-24 |
| EWS:FLI1-fusion(ETS)/SK_N_MC-EWS:FLI1-ChIP-Seq(SRA014231)/Homer | 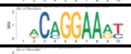 | 8324 | 47127 | 12770 | 83776 | 5.25E-22 | 1.15874492 | 2.54E-21 |
| Fli1(ETS)/CD8-FLI-ChIP-Seq(GSE20898)/Homer | 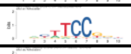 | 3371 | 52080 | 5524 | 91022 | 0.00438496 | 1.06654698 | 0.00520714 |
| GABPA(ETS)/Jurkat-GABPa-ChIP-Seq(GSE17954)/Homer | 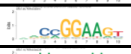 | 5120 | 50331 | 8469 | 88077 | 0.00248275 | 1.05794822 | 0.00314482 |
| IRF1(IRF)/PBMC-IRF1-ChIP-Seq(GSE43036)/Homer | 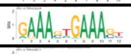 | 10345 | 45106 | 15949 | 80597 | 4.85E-26 | 1.15898673 | 9.22E-25 |
| IRF2(IRF)/Erythroblas-IRF2-ChIP-Seq(GSE36985)/Homer | 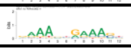 | 6897 | 48554 | 10886 | 85660 | 1.41E-11 | 1.11774643 | 3.35E-11 |

| | | | | | | | | |
|---|---|---|---|---|---|---|---|---|
| IRF4(IRF)/GM12878-IRF4-ChIP-Seq(GSE32465)/Homer | 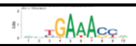 | 3829 | 51622 | 6024 | 90522 | 4.33E-07 | 1.1145977 | 6.58E-07 |
| ISRE(IRF)/ThioMac-LPS-Expression(GSE23622)/Homer | 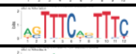 | 7882 | 47569 | 12031 | 84515 | 3.03E-22 | 1.16396813 | 1.92E-21 |
| PU.1-IRF(ETS:IRF)/Bcell-PU.1-ChIP-Seq(GSE21512)/Homer | 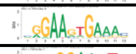 | 9939 | 45512 | 15695 | 80851 | 8.35E-17 | 1.12496493 | 2.89E-16 |
| PU.1(ETS)/ThioMac-PU.1-ChIP-Seq(GSE21512)/Homer | 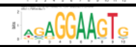 | 8538 | 46913 | 13665 | 82881 | 4.51E-11 | 1.10384232 | 9.53E-11 |
| SPDEF(ETS)/VCaP-SPDEF-ChIP-Seq(SRA014231)/Homer | 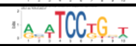 | 5820 | 49631 | 9344 | 87202 | 3.34E-07 | 1.09436589 | 5.28E-07 |
| T1ISRE(IRF)/ThioMac-Ifnb-Expression/Homer | 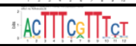 | 6430 | 49021 | 9597 | 86949 | 8.45E-24 | 1.1883746 | 6.42E-23 |
| TEAD(TEA)/Fibroblast-PU.1-ChIP-Seq(Unpublished)/Homer | 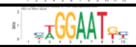 | 5583 | 49868 | 8307 | 88239 | 2.74E-21 | 1.18920701 | 1.16E-20 |

| Table S7. IRF8 motif enrichment in IRF8 KO plasmablasts | | | | | | | | |
|---|---|---|---|---|---|---|---|---|
| Motif | motif DNA sequence logo | a | b | c | d | pvalue | odds | FDR |
| MA0051.1 | | 932 | 11394 | 10983 | 128688 | 0.23463208 | 0.95841799 | 0.33022293 |
| MA0652.1 | | 598 | 11728 | 7298 | 132373 | 0.07531782 | 0.92484562 | 0.11925321 |
| MA0653.1 | | 939 | 11387 | 10999 | 128672 | 0.31950312 | 0.96468566 | 0.42398841 |
| MA0098.3 | 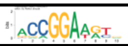 | 800 | 11526 | 7681 | 131990 | 6.63E-06 | 1.19270722 | 1.57E-05 |
| MA0772.1 | | 1139 | 11187 | 13619 | 126052 | 0.06802052 | 0.94234449 | 0.11620458 |
| MA1418.1 | 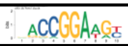 | 1416 | 10910 | 16149 | 123522 | 0.81411897 | 0.9927436 | 0.8593478 |
| MA1419.1 | | 549 | 11777 | 6579 | 133092 | 0.2052253 | 0.94303526 | 0.29994467 |
| MA1420.1 | | 464 | 11862 | 5283 | 134388 | 0.94109673 | 0.99503863 | 0.94109673 |
| MA1484.1 | | 771 | 11555 | 6924 | 132747 | 9.71E-10 | 1.27923566 | 1.23E-08 |
| MA1509.1 | | 135 | 12191 | 1400 | 138271 | 0.3235701 | 1.0936693 | 0.42398841 |
| AP-1(bZIP)/ThioMac-PU.1-ChIP-Seq(GSE21512)/Homer | | 1249 | 11077 | 15194 | 124477 | 0.01057357 | 0.92373402 | 0.01953399 |
| bZIP:IRF(bZIP,IRF)/Th17-BatF-ChIP-Seq(GSE39756)/Homer | | 1663 | 10663 | 18582 | 121089 | 0.55216919 | 1.01630829 | 0.60912271 |
| E2A(bHLH),near_PU.1/Bcell-PU.1-ChIP-Seq(GSE21512)/Homer | | 1272 | 11054 | 13823 | 125848 | 0.1355922 | 1.0476481 | 0.20610014 |
| EHF(ETS)/LoVo-EHF-ChIP-Seq(GSE49402)/Homer | 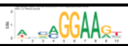 | 1748 | 10578 | 17648 | 122023 | 1.18E-06 | 1.14253421 | 3.44E-06 |
| ELF1(ETS)/Jurkat-ELF1-ChIP-Seq(SRA014231)/Homer | 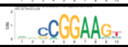 | 755 | 11571 | 6897 | 132774 | 1.98E-08 | 1.2561105 | 9.41E-08 |
| ELF5(ETS)/T47D-ELF5-ChIP-Seq(GSE30407)/Homer | 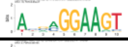 | 1263 | 11063 | 12337 | 127334 | 2.33E-07 | 1.17830733 | 8.85E-07 |
| Elk1(ETS)/Hela-Elk1-ChIP-Seq(GSE31477)/Homer | 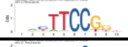 | 578 | 11748 | 5486 | 134185 | 4.94E-05 | 1.20340331 | 0.00010422 |
| Elk4(ETS)/Hela-Elk4-ChIP-Seq(GSE31477)/Homer | 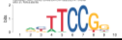 | 511 | 11815 | 4815 | 134856 | 8.23E-05 | 1.21132379 | 0.00016457 |
| ERG(ETS)/VCaP-ERG-ChIP-Seq(GSE14097)/Homer | 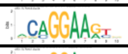 | 2217 | 10109 | 22342 | 117329 | 1.33E-08 | 1.15173826 | 8.43E-08 |
| ETS1(ETS)/Jurkat-ETS1-ChIP-Seq(GSE17954)/Homer | 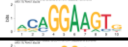 | 1938 | 10388 | 18967 | 120704 | 7.52E-11 | 1.18722224 | 1.43E-09 |
| Ets1-distal(ETS)/CD4+-PolII-ChIP-Seq(Barski et al.)/Homer | 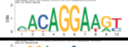 | 2148 | 10178 | 21437 | 118234 | 1.71E-09 | 1.16403195 | 1.63E-08 |
| ETS:E-box(ETS,bHLH)/HPC7-Scl-ChIP-Seq(GSE22178)/Homer | 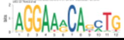 | 1705 | 10621 | 18187 | 121484 | 0.0107951 | 1.07231648 | 0.01953399 |
| ETS(ETS)/Promoter/Homer | 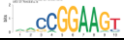 | 1001 | 11325 | 9097 | 130574 | 2.32E-11 | 1.26868283 | 8.81E-10 |
| ETS:RUNX(ETS,Runt)/Jurkat-RUNX1-ChIP-Seq(GSE17954)/Homer | 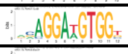 | 1757 | 10569 | 17825 | 121846 | 2.80E-06 | 1.13640769 | 7.10E-06 |
| ETV1(ETS)/GIST48-ETV1-ChIP-Seq(GSE22441)/Homer | 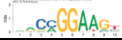 | 1541 | 10785 | 15309 | 124362 | 2.62E-07 | 1.160679 | 9.04E-07 |
| EWS:ERG-fusion(ETS)/CADO_ES1-EWS:ERG-ChIP-Seq(SRA014231)/Homer | 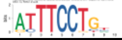 | 2108 | 10218 | 21443 | 118228 | 3.81E-07 | 1.13750083 | 1.21E-06 |
| EWS:FLI1-fusion(ETS)/SK_N_MC-EWS:FLI1-ChIP-Seq(SRA014231)/Homer | 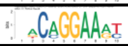 | 1931 | 10395 | 19163 | 120508 | 3.63E-09 | 1.16814315 | 2.76E-08 |
| Fli1(ETS)/CD8-FLI-ChIP-Seq(GSE20898)/Homer | 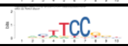 | 855 | 11471 | 8040 | 131631 | 1.72E-07 | 1.22029851 | 7.25E-07 |
| GABPA(ETS)/Jurkat-GABPa-ChIP-Seq(GSE17954)/Homer | 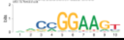 | 1276 | 11050 | 12313 | 127358 | 1.93E-08 | 1.19438752 | 9.41E-08 |
| IRF1(IRF)/PBMC-IRF1-ChIP-Seq(GSE43036)/Homer | | 2156 | 10170 | 24138 | 115533 | 0.55935871 | 1.01468864 | 0.60912271 |
| IRF2(IRF)/Erythroblas-IRF2-ChIP-Seq(GSE36985)/Homer | | 1469 | 10857 | 16314 | 123357 | 0.42993633 | 1.02309551 | 0.52701873 |

| | | | | | | | | |
|---|---|---|---|---|---|---|---|---|
| IRF4(IRF)/GM12878-IRF4-ChIP-Seq(GSE32465)/Homer | | 824 | 11502 | 9029 | 130642 | 0.34003751 | 1.03657307 | 0.43071418 |
| ISRE(IRF)/ThioMac-LPS-Expression(GSE23622)/Homer | | 1621 | 10705 | 18292 | 121379 | 0.86729836 | 1.00479793 | 0.89073886 |
| PU.1-IRF(ETS:IRF)/Bcell-PU.1-ChIP-Seq(GSE21512)/Homer | | 2103 | 10223 | 23531 | 116140 | 0.54699853 | 1.01531921 | 0.60912271 |
| PU.1(ETS)/ThioMac-PU.1-ChIP-Seq(GSE21512)/Homer | 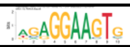 | 1981 | 10345 | 20222 | 119449 | 2.04E-06 | 1.13116176 | 5.53E-06 |
| SPDEF(ETS)/VCaP-SPDEF-ChIP-Seq(SRA014231)/Homer | 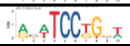 | 1368 | 10958 | 13796 | 125875 | 1.86E-05 | 1.13901393 | 4.16E-05 |
| T1ISRE(IRF)/ThioMac-Ifnb-Expression/Homer | | 1280 | 11046 | 14747 | 124924 | 0.56103407 | 0.9816272 | 0.60912271 |
| TEAD(TEA)/Fibroblast-PU.1-ChIP-Seq(Unpublished)/Homer | | 1182 | 11144 | 12708 | 126963 | 0.07033435 | 1.05969814 | 0.11620458 |
